# Quantitative Susceptibility Mapping for Differentiating Hydroxyapatite and Calcium Oxalate Breast Calcifications at 3T: A Phantom Study

**DOI:** 10.64898/2026.07.15.738683

**Authors:** Klara Mišak, Enrico De Vita, Chris A Clark, Matt T Cashmore, Simon Walker-Samuel

## Abstract

**Objective:** Breast microcalcifications cannot be differentiated by composition on mammography, and 70 to 80% of those biopsied prove benign. Type I calcifications are calcium oxalate (CaOx) and associate with benign processes; Type II are hydroxyapatite (HA) and correlate with malignancy. This work assesses whether quantitative susceptibility mapping (QSM) can distinguish the two at 3 T, and how much of the susceptibility present it recovers.

**Approach:** HA and CaOx granules were co-embedded in alginate phantoms in three formulations: undoped, adipose-matched and fibroglandular-matched. Four replicates per condition were imaged at 0.70 and 0.86 mm isotropic resolution. The reconstruction pipeline, including the choice of background field removal, was verified stage by stage against a digital twin of known susceptibility. R2* was evaluated from the same acquisition.

**Main results:** In undoped alginate, HA was detected in all eight measurements and CaOx in none. Recovery of known susceptibility was 18.2%, lost mainly to partial volume rather than reconstruction. Simulation places the susceptibility of the CaOx granules below 0.05 ppm relative to the gel. R2* was elevated at both particle types, 4.5 times more for HA. Fifteen of 48 measurements were affected by air retained within the porous granules, which can invert the sign of the recovered susceptibility.

**Significance:** In undoped alginate, susceptibility detects HA alone while R2* is elevated at both, so together they distinguish HA, CaOx and no particle from one acquisition. In simulation, the 50% detection diameter rises from 0.40 mm at 0.70 mm resolution to 1.07 mm at 1.50 mm, against a clinical range of 0.1 to 1.0 mm. Trapped air is detectable from the sign of the local field without reconstruction.

## 1. Introduction

Breast cancer remains the most diagnosed cancer among women worldwide, with survival rates strongly dependent on the stage at detection (Akram et al., 2017; Winters et al., 2017). Microcalcifications are microscopic calcium deposits typically 0.1 to 1.0 mm in diameter and serve as critical early indicators of malignancy, accounting for 85 to 95% of ductal carcinoma in situ (DCIS) detections (Baker et al., 2010; Henrot et al., 2014; Logullo et al., 2022). Mammography is highly effective at visualising these deposits. However, biopsies prompted by microcalcifications prove benign in 70 to 80% of cases (Bent et al., 2012). Reducing unnecessary biopsies while maintaining cancer detection sensitivity therefore represents a pressing clinical need.

The underlying limitation is the inability of current imaging modalities to differentiate calcifications by chemical composition. Breast calcifications fall into two categories (Frappart et al., 1984, 1986): Type I, composed of calcium oxalate dihydrate (CaOx, CaC₂O₄·2H₂O), associate primarily with benign secretory processes, whereas Type II, comprising hydroxyapatite (HA, Ca₁₀(PO₄) ₆(OH)₂), correlate strongly with malignancy. Mixed presentations can occur, though the dominant mineral type retains diagnostic significance (Bonfiglio et al., 2018; Henrot et al., 2014).

Both minerals are diamagnetic, and the susceptibility difference between them is the physical basis for differentiation. Susceptibilities recovered from biological calcium deposits are of the order of 1 part per million (ppm) relative to the surrounding medium, so the measurement must resolve a difference smaller than that.

Quantitative susceptibility mapping (QSM) exploits this contrast by solving the inverse field-to-source problem from multi-echo gradient echo phase data, yielding direct quantification of tissue magnetic susceptibility in ppm (Deistung et al., 2017). Unlike susceptibility-weighted imaging (SWI), which provides only qualitative contrast through T₂*-weighted signal loss and filtered phase (Haacke et al., 2004), QSM enables compositional characterisation and could distinguish HA from CaOx non-invasively. R2* relaxation rate mapping, derived from the same multi-echo magnitude data, offers an independent contrast mechanism sensitive to local field inhomogeneity and obtained without dipole inversion (Wang & Liu, 2015).

Compositional differentiation has been demonstrated in phantoms by two distinct approaches. At 9.4 T, CaOx and HA crystals of 200 to 500 µm were embedded in agarose and the apparent crystal area on gradient echo images compared against that on T₂-weighted images as a function of echo time (Mustafi et al., 2015); blooming artefact extent was more than twice as large for HA as for CaOx. The measurement is of artefact extent rather than of susceptibility and does not involve solution of the inverse problem. More recently, an ex vivo study of 53 renal calculi at 1.5 and 3 T reported −1.25 ± 0.16 ppm for carbonate apatite and −1.08 ± 0.13 ppm for CaOx (Mücke et al., 2026). Stones below 3.0 mm in diameter were excluded from that analysis owing to partial volume effects (PVE), and the reported values derive from stones of 3 to 10 mm at voxel sizes of 0.94 to 1.0 mm isotropic.

Neither approach establishes whether the two mineral types can be distinguished by quantified susceptibility at clinical field strength when the particle diameter is comparable to the acquisition voxel size, which is the regime in which breast microcalcifications are imaged. That is the question addressed here. Susceptibility is quantified in ppm rather than inferred from artefact extent, at 3 T rather than ultra-high field, for particles of 1 to 2 mm at 0.70 and 0.86 mm isotropic resolution: a regime in which PVE is not a nuisance to be excluded, but the dominant determinant of the measurement.

Existing breast MRI phantoms address quality assurance rather than calcification composition, as do the standard system phantoms (Keenan et al., 2016, 2021, 2024; Stupic et al., 2021). Phantoms with HA and CaOx particles co-embedded in a single gel volume were therefore developed, in three formulations spanning breast relaxation times, with four replicates per condition.

QSM pipelines have largely been developed for neuroimaging, where sources are spatially distributed across well-defined geometry (Bilgic et al., 2024). The sources here are focal and bounded by air, and background field removal must operate close to them. No stage of the reconstruction could therefore be assumed to transfer. Each was verified against a digital twin phantom with known ground truth, following the practice established by the QSM Reconstruction Challenge (Marques et al., 2021). Background field removal was chosen by the same comparison rather than by convention. Data were acquired at two isotropic resolutions to test whether PVE rather than susceptibility contrast sets the detection limit. R2* was evaluated from the same magnitude data as an independent contrast, and the two assessed together as a two-parameter scheme.

## 2. Methods

### 2.1 Synthetic Calcification Particle Fabrication

Synthetic calcification particles representing Type I (CaOx) and Type II (HA) breast calcifications were fabricated by reproducing a compression granulation method (Ghammraoui et al., 2021). Reagent-grade HA and CaOx dihydrate powders (>99% purity; Sigma-Aldrich, St Louis, MO, USA) were used as base materials. For each mineral type, 5 g of powder was mixed with 50 mg polyvinylpyrrolidone K30 (PVP K30; 1% w/w) and bound with 3% PVP K30 in ethanol (powder-to-binder ratio 5:1). The wet paste was vacuum-dried at 30 °C for 48 hours, then compressed using an arbour press for 60 seconds.

Compressed tablets were crushed and size-selected by sequential sieving through 2.0, 1.0 and 0.5 mm aperture meshes. The 1.0–2.0 mm fraction was used for all primary experiments, yielding irregularly shaped particles with a mean long axis of 2.02 ± 0.62 mm and short axis of 1.35 ± 0.25 mm.

As the particles are compression granules rather than solid crystals, their effective susceptibility is diluted by internal porosity and by gel infiltration of the pore space. Packing fraction was not measured here; the solid fractions used are taken from the granule densities reported for this compression protocol (Ghammraoui et al., 2021).

### 2.2 Gel Formulation

Alginate hydrogels were prepared using a standardised protocol (Liang et al., 2026). All formulations used 2% (w/v) sodium alginate dissolved in 1.17% (w/v) NaCl saline solution (Lin et al., 2025).

Three gel formulations were prepared. Pure crosslinked alginate (no paramagnetic dopants) served as a susceptibility reference baseline. Two relaxometry-matched formulations were additionally prepared by incorporating nickel chloride (NiCl₂) for T1 shortening and manganese chloride (MnCl₂) for T2 shortening (Boretius & Frahm, 2011; Clarkson et al., 2025), with dopant concentrations calculated from published relaxivity equations at 3 T (Boss et al., 2018) to approximate human breast tissue relaxation times (Rakow-Penner et al., 2006) (**Table 1**).

**Table 1.** Gel formulations. All formulations used 2% (w/v) sodium alginate in 1.17% (w/v) NaCl. Target relaxation times are the breast tissue values reported by (Rakow-Penner et al., 2006). Dopant Δχ is the calculated Curie-law contribution (Coey, 2010) to the bulk susceptibility at 20 °C, relative to undoped alginate. Dashes indicate not applicable.

| Formulation | NiCl<sub>2</sub><br>(mM) | MnCl<sub>2</sub><br>(mM) | Target T<sub>1</sub><br>(ms) | Target T<sub>2</sub><br>(ms) | Dopant<br>$\Delta\chi$<br>(ppm) |
| --- | --- | --- | --- | --- | --- |
| Pure alginate | — | — | — | — | — |
| FGT-matched | 0.42 | 1.14 | 1200–1680 | 40–60 | +0.24 |
| Adipose-matched | 3.10 | 1.13 | 350–450 | 50–70 | +0.38 |

Each gel formulation was centrifuged (3000 rpm, 10 min) to remove trapped air from the gel.

### 2.3 Phantom Assembly and Experimental Design

Each phantom consisted of a 50 mL conical centrifuge tube (Falcon; Corning Inc., Corning, NY, USA) containing one HA and one CaOx calcification particle embedded within the same alginate gel volume, separated by approximately 15–20 mm along the tube axis. This combined-tube design ensured that both mineral types shared identical imaging conditions, including B₀ shimming, QSM zero-point reference, and gel baseline susceptibility, enabling direct within-tube comparison without inter-phantom baseline confounds.

Phantom assembly followed a controlled partial-gelation embedding protocol. A base gel layer (10 mL alginate solution + 2 mL crosslinking solution) was dispensed into the tube and allowed to partially gel for approximately 8–10 minutes until firm enough to support a particle. The HA particle was placed at a defined axial position, an intermediate gel layer (5 mL alginate + 1 mL crosslinking solution) was added after 3 minutes to allow the microcalcification to embed into the gel, and after a further 10–15 minutes of partial gelation the CaOx particle was placed at the target separation distance. A final encapsulation layer (10 mL alginate + 2 mL crosslinking solution) was applied (**Fig. 1**).

**Fig. 1.**
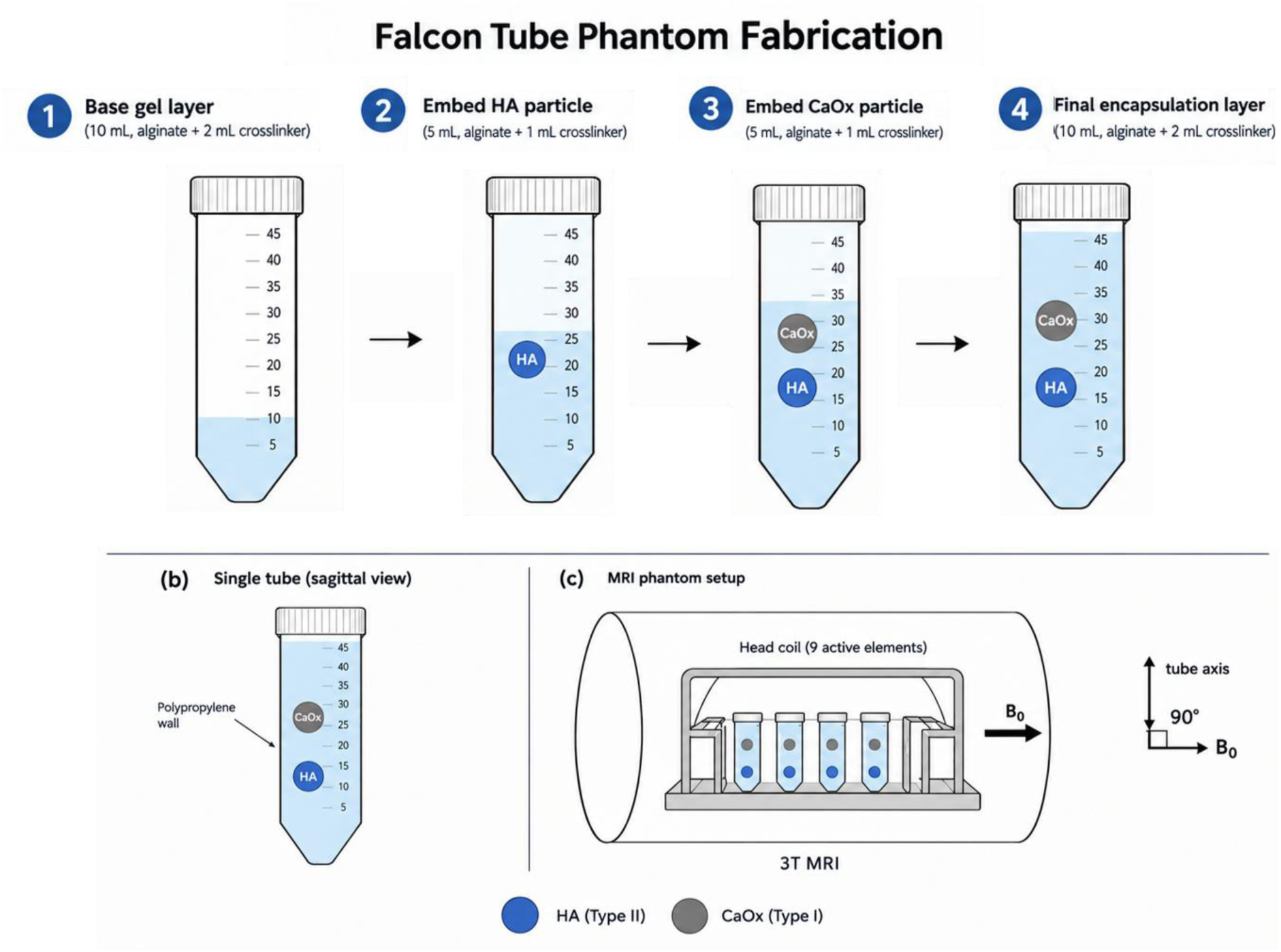
Phantom fabrication and experimental setup. (Top) Five-step partial-gelation embedding protocol for co-embedding HA and CaOx particles within a single 50 mL Falcon tube, with gel volumes and crosslinker ratios indicated. (a) Sagittal view of a completed tube showing HA and CaOx particles separated by 15–20 mm within BaCl₂-crosslinked alginate gel. (b) MRI phantom setup: four tubes positioned in a 64-channel head coil at 3T with tube axes aligned perpendicular to B₀.

All phantoms were crosslinked using 0.1 M barium chloride (BaCl₂) solution. Calcium chloride (CaCl₂), the standard alginate crosslinker (Liang et al., 2026), was not used because preliminary experiments demonstrated that the crosslinking solution progressively dissolved CaOx particles, with visible particle degradation within 10 minutes. This is attributed to disruption of the CaOx crystal lattice by excess Ca²⁺ ions in solution. Barium ions crosslink alginate through the same egg-box coordination mechanism as calcium but do not participate in oxalate chelation, preserving particle integrity across both mineral types (Mørch et al., 2012).

Following assembly, tubes equilibrated to scanner room temperature (19–20 °C) for a minimum of 24 hours prior to imaging. Twelve phantoms were fabricated in total: four replicates per gel type (pure alginate, adipose relaxometry-matched, FGT relaxometry-matched).

### 2.4 MRI Acquisition

All imaging was performed on a 3T Siemens MAGNETOM Prisma scanner using the manufacturer’s 64-channel head coil. Multi-echo gradient echo (ME-GRE) data were acquired at two isotropic resolutions (0.70 mm and 0.86 mm) with seven echoes per acquisition (**Table 2**).

**Table 2.**
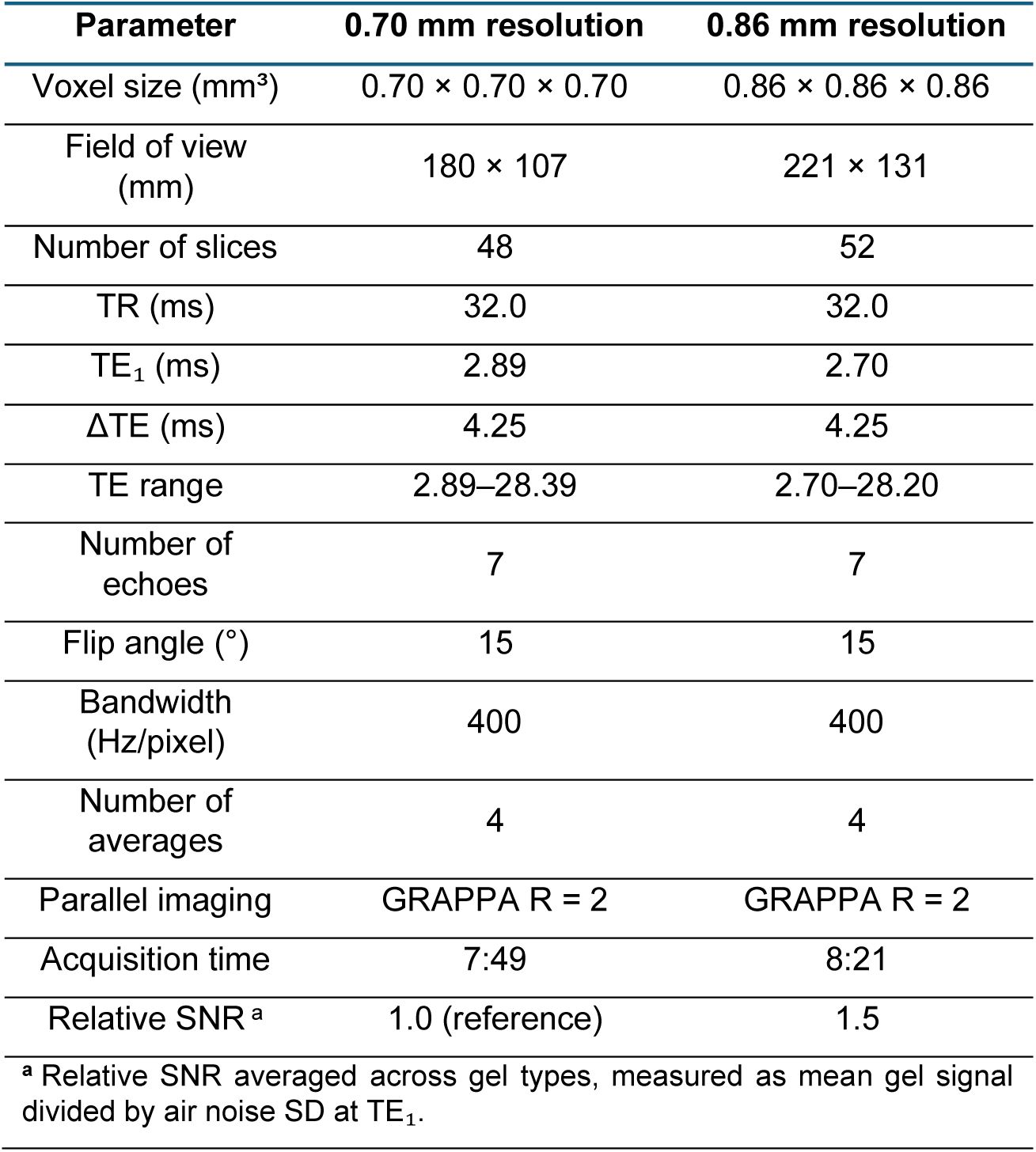
Acquisition parameters for multi-echo gradient echo sequences used for QSM. All acquisitions performed at 3T (2.895T, Siemens MAGNETOM Prisma) using a 64-channel head coil.

Four phantoms were positioned in the coil per scan session with the tube long axes perpendicular to B₀.

The field strength was 2.895 T (Larmor frequency 123.244417 MHz); this value rather than the nominal 3 T was used throughout the reconstruction.

### 2.5 QSM Processing Pipeline

A QSM pipeline was implemented in Python (3.10, NumPy 1.24, SciPy 1.11) following the ISMRM QSM Consensus recommendations (Bilgic et al., 2024) (**Fig. 2**).

**Fig. 2.**
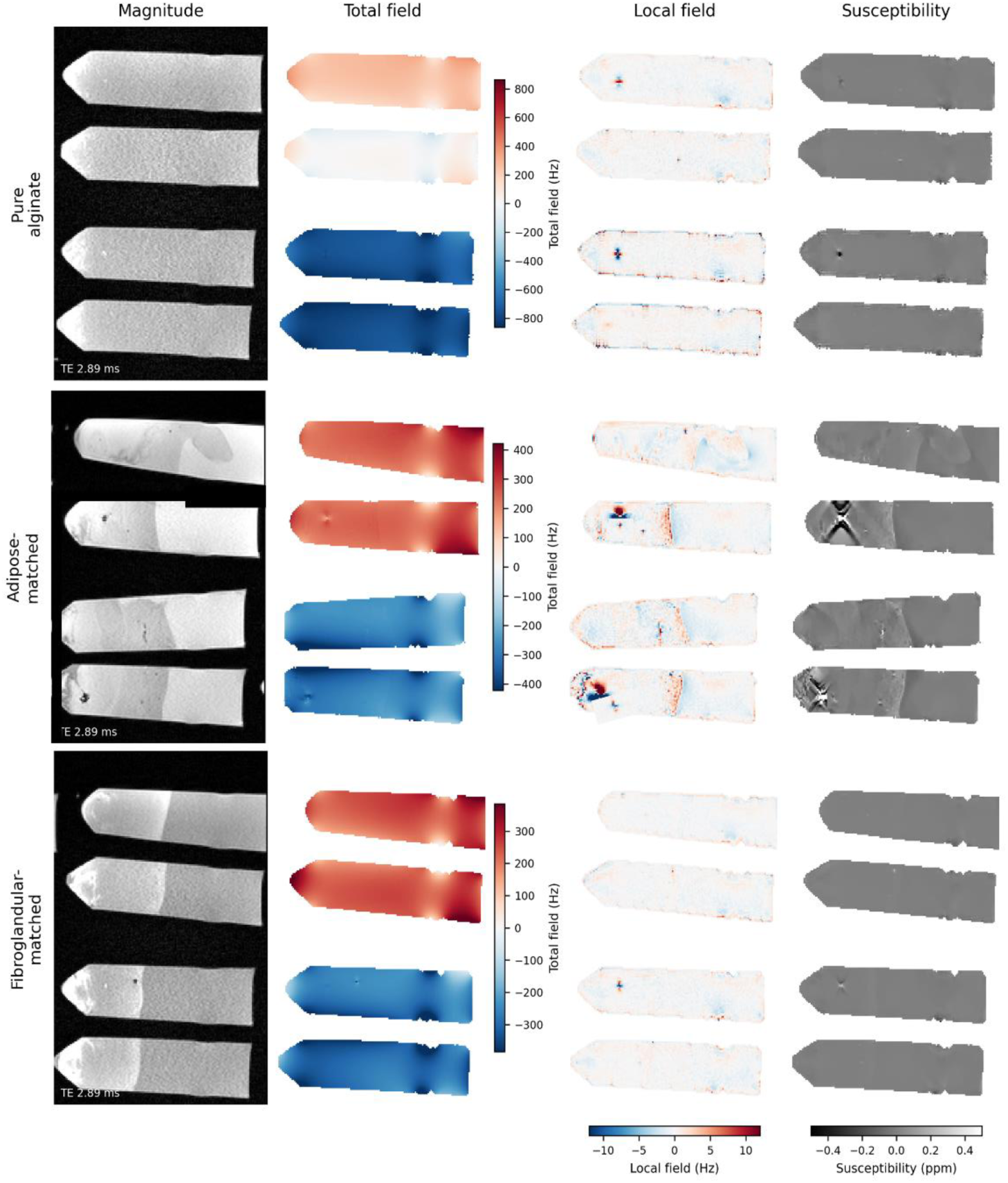
Reconstruction stages for each gel formulation at 0.70 mm isotropic resolution, one sagittal slice per formulation with B₀ vertical. Left to right: magnitude at the first echo (TE 2.89 ms), total field, local field after background field removal by projection onto dipole fields, and susceptibility. Tubes are numbered from the top; each contains one HA and one CaOx particle, though not all particles lie within the displayed slice. The local field and susceptibility columns share a common window across rows. The total field window is set per row from the 1st to 99th percentile of the displayed slice, since inter-tube field offsets differ between formulations. Boundaries visible in the adipose-matched magnitude images correspond to the partial-gelation embedding layers (**Section 2.3**). Streaking is visible in the susceptibility maps at the adipose-matched particle locations, in rows 2 and 4 of that group, and is discussed in **Section 3.3**; the bipolar local field features at the same locations are the signature on which the air criterion of **Section 2.9** operates.

Available QSM toolboxes are built around neuroimaging workflows, including Brain Imaging Data Structure (BIDS) organisation and brain-extraction masking, and required substantial modification for isolated tubes. Verification was the more substantial constraint: comparing each stage against a known result requires access to intermediate quantities that packaged implementations do not expose. The reconstruction was therefore implemented with explicit per-stage control and verified as described in **Section 2.6**, using established implementations of individual stages (**Sections 2.5.2, 2.5.3**).

#### 2.5.1 Masking

Binary tissue masks were generated by thresholding the mean magnitude image at 10% of maximum, followed by phase-quality refinement (voxels with temporal coherence below 0.3 excluded) and then hole-filling. The order matters: localised dephasing around the calcifications reduces temporal coherence at precisely the locations of interest, so refinement removes the particles from the mask, and filling afterwards recovers them (Stewart et al., 2022).

A two-voxel morphological erosion was then applied to remove the low SNR at the gel–tube interface, where residual phase-unwrapping errors were found to concentrate (**Section 2.5.2**).

#### 2.5.2 Phase unwrapping

Phase unwrapping used ROMEO with 4D template unwrapping across all seven echoes. (Dymerska et al., 2021).

Whole-volume unwrapping connects physically disconnected tubes along a minimum spanning tree passing through air and can fail when inter-tube field offsets are large. The magnitude of the offset was found not to predict failure: one dataset with a 720 Hz spread across tubes gave 15,030 residual errors, while another with a 1030 Hz spread gave none.

A measured criterion was therefore used. Residual errors were quantified after each reconstruction by counting voxel-to-voxel field differences within 15% of the aliasing frequency (1/ΔTE = 235.3 Hz at 0.70 mm), and where this count exceeded 100, unwrapping was repeated independently for each tube on a cropped volume. Four of six datasets required per-tube unwrapping.

Residual errors, where present, were localised: all lay within 2 mm of the tube wall (median 0.70 mm), in voxels with magnitude 29% of the gel mean, and none within 5 mm of any calcification.

#### 2.5.3 Field mapping and background field removal

Total field maps were computed by magnitude-weighted linear regression of unwrapped phase against echo time, with weights w(TE) = |S(TE)|².

The total field is dominated by contributions from sources outside the region of interest (ROI), principally the susceptibility difference at the surrounding air. Background field removal used projection onto dipole fields (PDF) (T. Liu, Khalidov, de Rochefort, et al., 2011), which models these contributions as arising from dipole sources outside the mask and subtracts the field they would produce.

PDF was selected after direct comparison against V-SHARP (Özbay et al., 2017) and Laplacian boundary value (LBV) (Zhou et al., 2014) methods, applied to identical field maps with identical dipole inversion and evaluated against known susceptibility in the digital twin phantom (**Section 2.6**). The comparison and the basis for the choice are reported in **Section 3.2**.

#### 2.5.4 Dipole inversion

Susceptibility maps were reconstructed by total variation regularised dipole inversion, solved by the alternating direction method of multipliers (ADMM), minimising

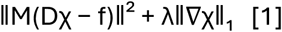

where M is the mask, D the dipole kernel, f the local field and λ the regularisation weight. Total variation was chosen to suppress streaking while preserving the sharp edges of a compact source in uniform gel, on which the localisation required by the detection analysis depends.

The χ-subproblem was solved by preconditioned conjugate gradient rather than by direct division in k-space. The data-consistency operator DᴴMD is not diagonal in k-space when a mask is present, so k-space division solves the unmasked problem instead. With the unmasked solve, recovered susceptibility depended on the ADMM penalty parameter μ, which should affect only the rate of convergence. With the masked operator solved by conjugate gradient, recovered susceptibility varied by 0.6% across a tenfold range of μ, confirming convergence to the intended solution. Reconstruction used 60 ADMM iterations with a thresholded k-space division warm start (threshold 0.2).

The regularisation weight was selected by minimising the error in recovered mean susceptibility against known susceptibility in the digital twin phantom (**Section 2.6**), giving λ = 5 × 10⁻⁴. An L-curve criterion was not usable here: the log–log curve of data fidelity against total variation showed no corner over the range tested, from 5 × 10⁻⁵ to 5 × 10⁻³, with an approximately constant slope of −0.9 below 10⁻³, so corner-finding returns an endpoint rather than an optimum. Selection by whole-tube root-mean-square error favoured a slightly larger value, reflecting a trade-off between smoothness of the gel and fidelity of the source (**Supplementary Table S1**). The inversion is also the stage at which this pipeline differs most from published calcification reconstructions, so detection outcomes were compared against morphology-enabled dipole inversion (MEDI) (J. Liu et al., 2012) on the same local field maps (**Section 3.2**).

### 2.6 Pipeline Validation

Reconstruction was verified against a digital twin: a numerical phantom of known susceptibility, processed through the same pipeline as the experimental data, so that recovered values could be compared against the values supplied. Recovery is reported against one of two references. Recovery against input is the recovered value as a percentage of the input susceptibility and includes PVE dilution. Recovery against partial-volume ground truth is the recovered value as a percentage of the input susceptibility averaged over the same ROI and isolates the reconstruction. Both are computed on Δχ_mean_ unless stated otherwise.

#### 2.6.1 Digital twin construction

The twin reproduces the experimental geometry and acquisition. Four tubes of 27 mm inner diameter with 1.5 mm walls at −0.75 ppm were modelled with their long axes perpendicular to B₀, matching the scan orientation (**Section 2.4**). Gel susceptibility was set per formulation, taking undoped alginate as −9.03 ppm and adding the calculated dopant contributions of **Table 1**.

Mineral sources were modelled as randomly oriented ellipsoids with semi-axes drawn from the measured particle distribution (**Section 2.1**) and were constructed on a grid eight times finer than the acquisition voxel before averaging down, so that boundary voxels carry a volume-weighted mixture of particle and gel.

Multi-echo signal was synthesised at twice the acquisition resolution and the complex signal block-averaged to the acquisition grid. Within a voxel containing both gel and particle, sub-voxel magnetisation precesses at different rates, and the contributions add as vectors. Averaging susceptibility before synthesising signal replaces that sum with a scalar mean and removes the intravoxel phase cancellation.

R2* and signal-to-noise ratio were set per formulation to values measured from the acquired phantom data. Gaussian noise was added independently to the real and imaginary channels, giving Rician magnitude statistics, scaled by a further factor of 1.96 so that gel susceptibility variability in the simulation matched that measured in the phantoms. Receive sensitivity was applied as a profile extracted from the gel, whose homogeneity means that smooth spatial variation in its magnitude reflects the sensitivity profile; measured values ranged from 0.61 to 0.69 at the phantom edge relative to the centre. A residual shim gradient of ±300 Hz along B₀ was added.

The susceptibility distribution was zero-padded by 48 voxels before the dipole convolution. Fourier convolution treats the array as periodic, so without padding the adjacent periodic copies of the phantom interact and the internal field within the tubes is recovered approximately 55% low. With 48 voxels of padding, the residual deviation from the analytic infinite-cylinder value is 14.8%, which is the finite-length effect of the modelled tube rather than a numerical artefact.

The forward model was verified against analytic solutions for a sphere and a cylinder, and against the requirements of linearity and superposition (**Supplementary Table S2**).

#### 2.6.2 Detection sensitivity to particle size

The particles used experimentally were 1 to 2 mm, while clinical microcalcifications are 0.1 to 1.0 mm. The size at which detection fails was therefore established by simulation, at six equivalent diameters from 0.3 to 2.0 mm and three acquisition resolutions, of which only 0.70 mm was acquired experimentally. The detection criterion of **Section 2.8.3** was applied unchanged, with specificity assessed at 1000 particle-free locations. Solid mineral was simulated rather than the porous granules used experimentally, so the resulting diameters are a lower bound on the size required for detection. Baseline relaxation and noise were taken from the 0.70 mm pure alginate measurements, with signal-to-noise scaled with voxel volume as it would be at fixed acquisition time, and R2* elevation at the particle set to 175.5 s⁻¹, representative of the values measured in pure alginate (**Table 6**). Full parameters are given in **Supplementary Table S3**.

### 2.7 R2* Analysis

R2* maps were computed from the same multi-echo magnitude data. Mono-exponential decay was fitted voxel-wise by nonlinear least squares, with R2* constrained to 0–500 s⁻¹ and voxels failing to reach a coefficient of determination of 0.9 excluded.

Only echoes with SNR above 3 were included, with noise estimated from voxels outside the phantom at the final echo. Rician magnitude noise plateaus rather than decaying to zero, so fitting through the noise floor biases R2* downward, and the bias is larger for faster-decaying formulations. In practice the criterion excluded no echoes: the median number used was seven of seven in every dataset, and fitted rates were unchanged when all echoes were included unconditionally.

Mono-exponential fitting is appropriate here because the gels contain no lipid, so the magnitude decays monotonically (Hernando et al., 2013; Horng et al., 2013).

### 2.8 Region of Interest Definition and Statistical Analysis

#### 2.8.1 Localisation

Particle locations were identified on the magnitude images, in which the calcifications appear as signal voids. The magnitude is unaffected by dipole inversion, so the region is defined independently of the susceptibility map from which measurements are taken.

A spherical ROI of two-voxel radius was centred on each location, and a gel reference region placed in homogeneous gel within the same tube, at least 8 mm from any particle. All ROIs lay within the region in which background field removal produced valid output.

A fixed spherical region was used rather than segmenting the void on the magnitude images. The void is inflated by blooming, which scales with susceptibility, so HA and CaOx voids differ in extent (2.5 against 1.9 mm equivalent diameter in pure alginate at 0.70 mm) despite both mineral types being drawn from the same sieve fraction. Segmenting on magnitude would therefore give the two minerals systematically different region volumes, and the PVE dilution would differ between them.

#### 2.8.2 Metrics

All values are reported relative to a gel reference region placed in the same tube. Δχ_peak_ is the 5th percentile of susceptibility within the particle region minus the gel reference mean and is the primary measurement and the quantity to which the detection criterion is applied. Δχ_mean_, the mean over the particle region minus the same reference, is reported alongside it. R2* elevation is the 95th percentile within the particle region minus the gel reference mean.

The percentiles are chosen to sample where each effect is strongest. Diamagnetic susceptibility is concentrated at the particle core, while relaxation elevation concentrates in the voxels nearest the particle, where the intravoxel field gradient is steepest. With approximately 33 voxels in the region, the 5th percentile corresponds to the second most diamagnetic voxel, so Δχ_peak_ is only marginally more robust than a minimum and is reported as a detection statistic rather than as an unbiased estimate of the particle susceptibility.

#### 2.8.3 Detection criterion

A particle was classified as detected by susceptibility if both of the following held:

1. |Δχ_peak_| exceeded three times the standard deviation of the gel reference. The three-standard-deviation threshold follows the Currie limit-of-detection convention (Currie, 1999). The gel reference standard deviation was measured over a region of the same size as the particle region, since it depends strongly on region size: the whole-gel standard deviation was approximately five times that of a two-voxel-radius region.
2. A focal diamagnetic minimum was present, defined as the centre voxel being more diamagnetic than the mean of voxels at a radius of two voxels by at least one gel standard deviation. This distinguishes a localised susceptibility source from a diffuse offset or a noise excursion.

Detection by R2* was assessed separately, as ΔR2* exceeding three times the standard deviation of the gel reference R2*.

### 2.9 Identification of air-affected measurements

Both trapped air and calcium minerals produce magnitude signal voids, but they differ in the sign of their susceptibility: air is approximately +9.4 ppm relative to water, whereas both HA and CaOx are diamagnetic. A diamagnetic source produces a negative field lobe along B₀ and a positive lobe perpendicular to it; a paramagnetic source produces the reverse.

Measurements were classified on the sign of the local field along B₀, sampled in two regions of one-voxel radius placed 2.80 mm either side of each particle and averaged, referenced to gel in the same tube. The local field precedes dipole inversion, so the classification does not depend on the reconstruction or its regularisation.

The threshold was set empirically. The same sampling was applied at 2000 particle-free positions per condition, at least 10 mm from any particle and four voxels from the tube wall, and the 99.7th percentile of that distribution was taken as the threshold: +2.22 and +1.33 Hz for pure alginate at 0.70 and 0.86 mm, +2.83 and +2.93 for adipose-matched, and +1.91 and +1.52 for FGT-matched. These correspond to air pockets of 0.7 to 0.9 mm diameter, or 10 to 20% of the pore volume of a 2 mm granule, which is above the few per cent consistent with near-complete wetting and below anything that would materially dilute the mineral.

That null is sampled at least 10 mm from any particle, while the criterion is applied 2.80 mm from one, where the particle’s own field contributes. The CaOx locations in pure alginate provide a null at matched distance: eight measurements span −0.18 to +0.40 Hz with a standard deviation of 0.23 Hz, narrower than the particle-free null, so the thresholds above are conservative at that distance. These are the same locations whose non-detection places the CaOx bound in **Section 3.3.3**, so the two are not independent; the thresholds themselves are set from the particle-free null. Equivalent locations in the doped formulations are too few for an independent estimate, and the alginate standard deviation is used throughout.

Because HA produces its own negative lobe along B₀, air adjacent to an HA particle partially cancels while air adjacent to CaOx does not. The criterion is therefore more sensitive at CaOx locations, and the distribution of identifications between the two minerals should not be read as a difference in how much air each retained.

All measurements are reported. Air-affected locations are flagged in **Table 5** and **Fig. 5**, and their interpretation is addressed in **Sections 3.5 and 4**.

## 3. Results

### 3.1 Pipeline validation against ground truth

Recovery was quantified in the digital twin phantom across 80 independent measurements (five particle realisations × four noise realisations × four tubes) at an input susceptibility of −2.0 ppm, measured over a two-voxel-radius region of interest (ROI) to match the experimental analysis.

The pipeline recovered 18.2% of the input susceptibility, and the larger part of that loss was geometric rather than reconstructive (**Table 3**). A two-voxel-radius ROI contains approximately 33 voxels at 0.70 mm isotropic resolution, while a particle of the measured mean dimensions occupies approximately 12, so the measurement is diluted by the surrounding gel. Representative ground truth and reconstructed maps are shown in **Fig. 3**.

**Fig. 3.**
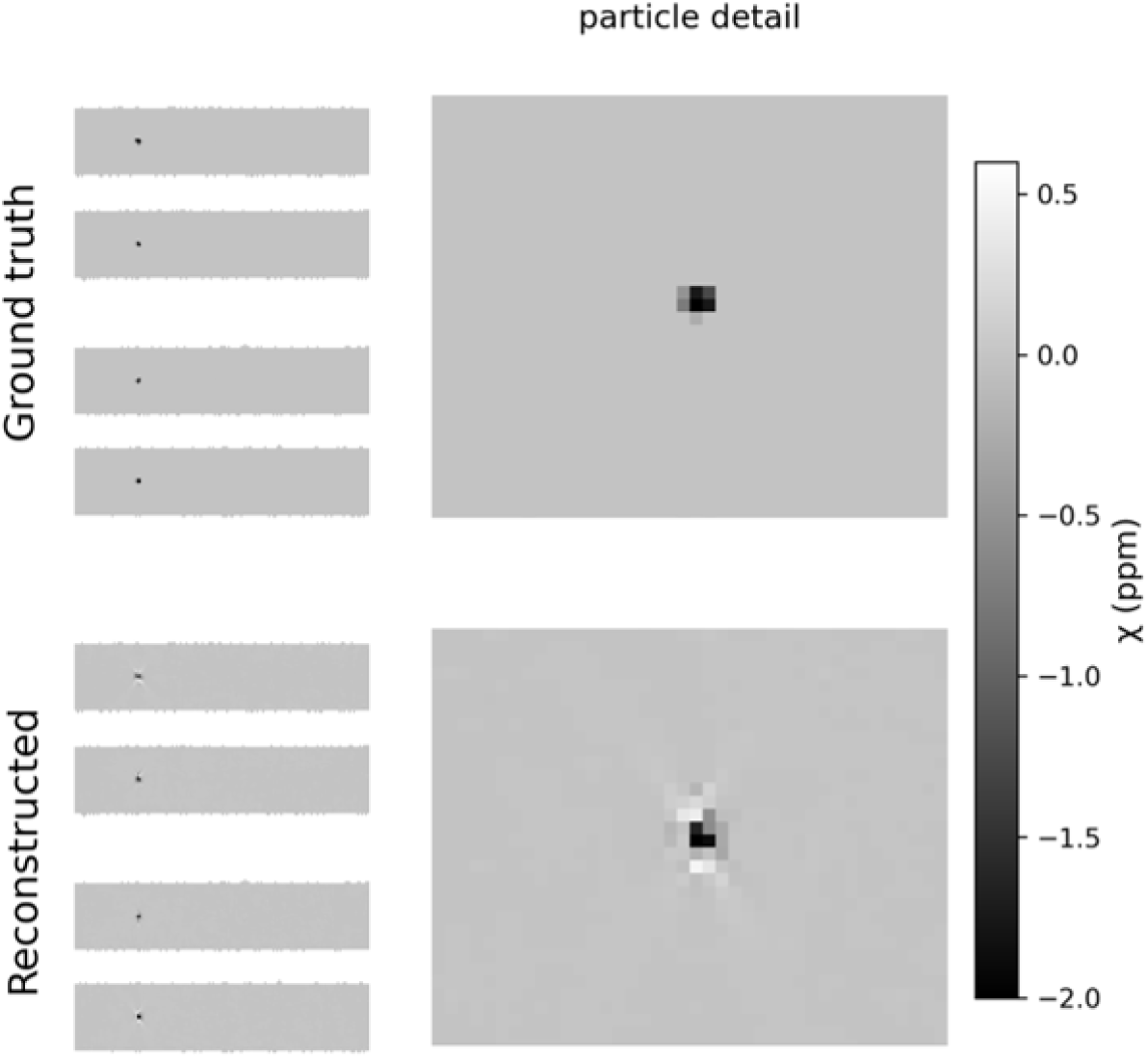
Digital twin susceptibility maps. (a) ground truth, the susceptibility distribution supplied to the forward model; (b) the corresponding reconstruction after the full pipeline. Right-hand panels show a magnified view of one HA particle. Both panels share a single colour scale. CaOx was modelled at −0.02 ppm, below the detection bound established in **Section 3.3.3**, and is correspondingly absent.

**Table 3.**
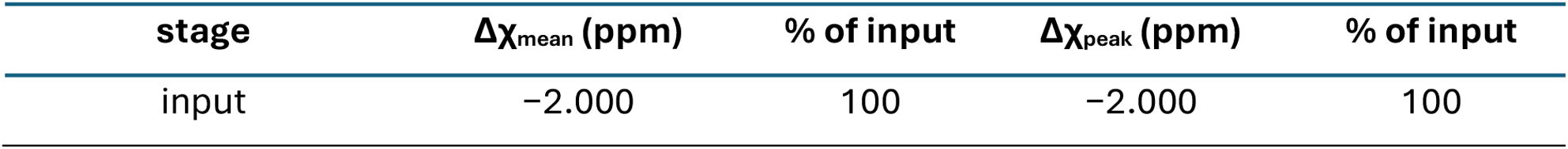
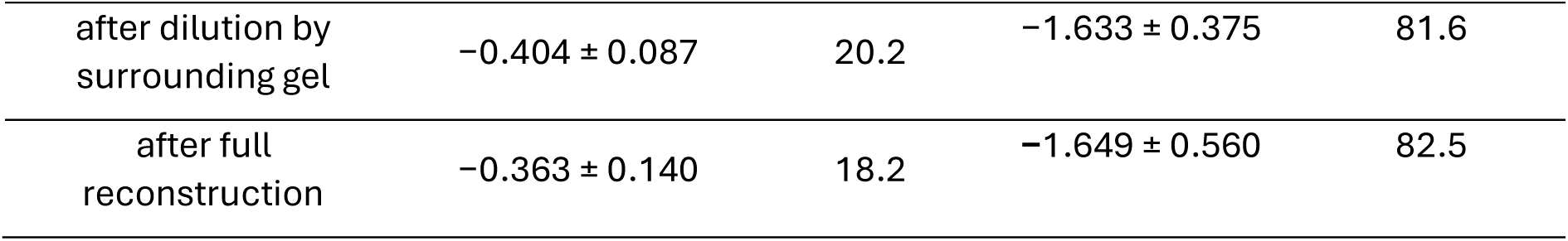
Susceptibility recovery in the digital twin phantom over a two-voxel-radius ROI, at an input susceptibility of −2.0 ppm. Values are mean ± standard deviation across 80 measurements. Percentages are recovery against input. PVE dilution applies to the region average; the 5th percentile samples the particle core and is diluted far less. Reconstruction retains 89.8% of the partial-volume ground truth for Δχmean and 101.0% for Δχpeak, the latter reflecting dipole ringing at the particle boundary; the percentile is reported as a detection statistic rather than as a quantitative estimate (**Section 2.8.2**).

Recovery was independent of the gel formulation, at 18.2%, 18.1% and 18.2% for pure alginate, adipose-matched and FGT-matched gels respectively, despite simulated baseline R2* spanning 5.9 to 84.2 s⁻¹ and signal-to-noise ratio spanning 27.3 to 44.0 across the three.

The two susceptibility metrics behave differently over the ROI. Δχ_peak_ recovered 82.5% of the input susceptibility against 18.2% for Δχ_mean_, since the percentile samples the particle core while the mean averages it with the surrounding gel. Their variability across the 80 measurements is comparable, with coefficients of variation of 34% and 39% respectively.

### 3.2 Comparison of reconstruction methods

The background field dominated the measurement: across the phantom datasets it accounted for 99.8% of the total field standard deviation, with a standard deviation of 454 Hz against approximately 1 Hz for the local field.

Three background field removal methods were applied to identical field maps with identical dipole inversion and compared against known ground truth in the digital twin phantom (**Table 4**). Comparison against ground truth is necessary here rather than merely convenient: on measured data, a method that smooths aggressively and one that removes background accurately both produce a low gel reference standard deviation, and the two cannot be distinguished without a known answer.

**Table 4.**
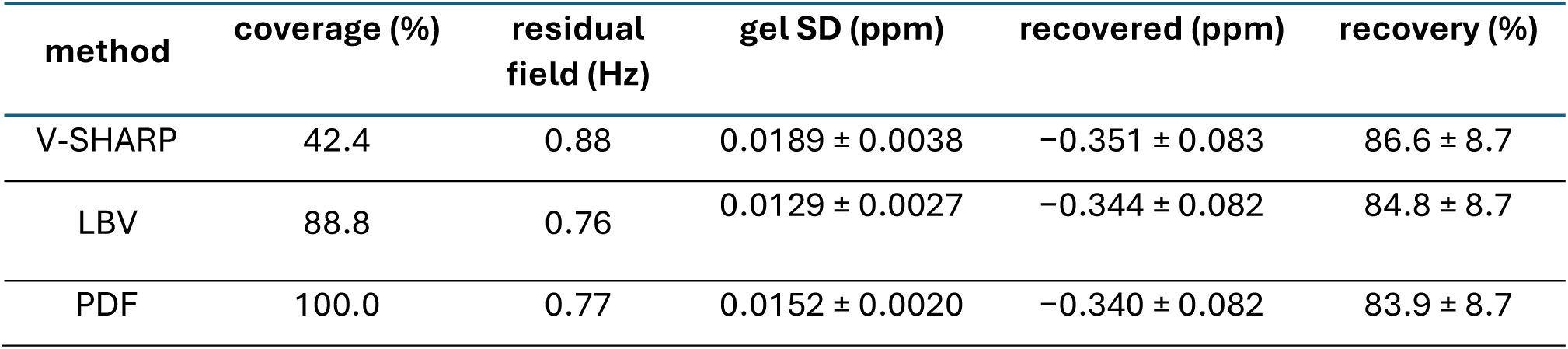
Background field removal methods compared in the digital twin, with identical dipole inversion. Coverage is the fraction of the masked volume with valid output; residual field is the local field SD 4 to 6 mm from the tube wall; gel SD is the susceptibility SD in the gel reference region. Recovered is Δχmean over a two-voxel-radius ROI, and recovery is against the partial-volume ground truth of −0.404 ± 0.087 ppm, as in Table 3. Values are mean ± standard deviation across five particle realisations, one noise realisation each, four tubes.

| method | coverage (%) | residual field (Hz) | gel SD (ppm) | recovered (ppm) | recovery (%) |
| --- | --- | --- | --- | --- | --- |
| V-SHARP | 42.4 | 0.88 | $0.0189 \pm 0.0038$ | $-0.351 \pm 0.083$ | $86.6 \pm 8.7$ |
| LBV | 88.8 | 0.76 | $0.0129 \pm 0.0027$ | $-0.344 \pm 0.082$ | $84.8 \pm 8.7$ |
| PDF | 100.0 | 0.77 | $0.0152 \pm 0.0020$ | $-0.340 \pm 0.082$ | $83.9 \pm 8.7$ |

All three recovered comparable susceptibility, within 2.9 percentage points of one another and well within the standard deviation across particle realisations. They differed substantially in coverage, the fraction of the masked volume for which each produced valid output (**Fig. 4**). Coverage rather than accuracy determined the choice: several ROIs fell outside the V-SHARP valid region in the experimental data, and a ROI that cannot be measured is of less use than one measured marginally less accurately. Projection onto dipole fields (PDF) was adopted for all subsequent analysis.

**Fig. 4.**
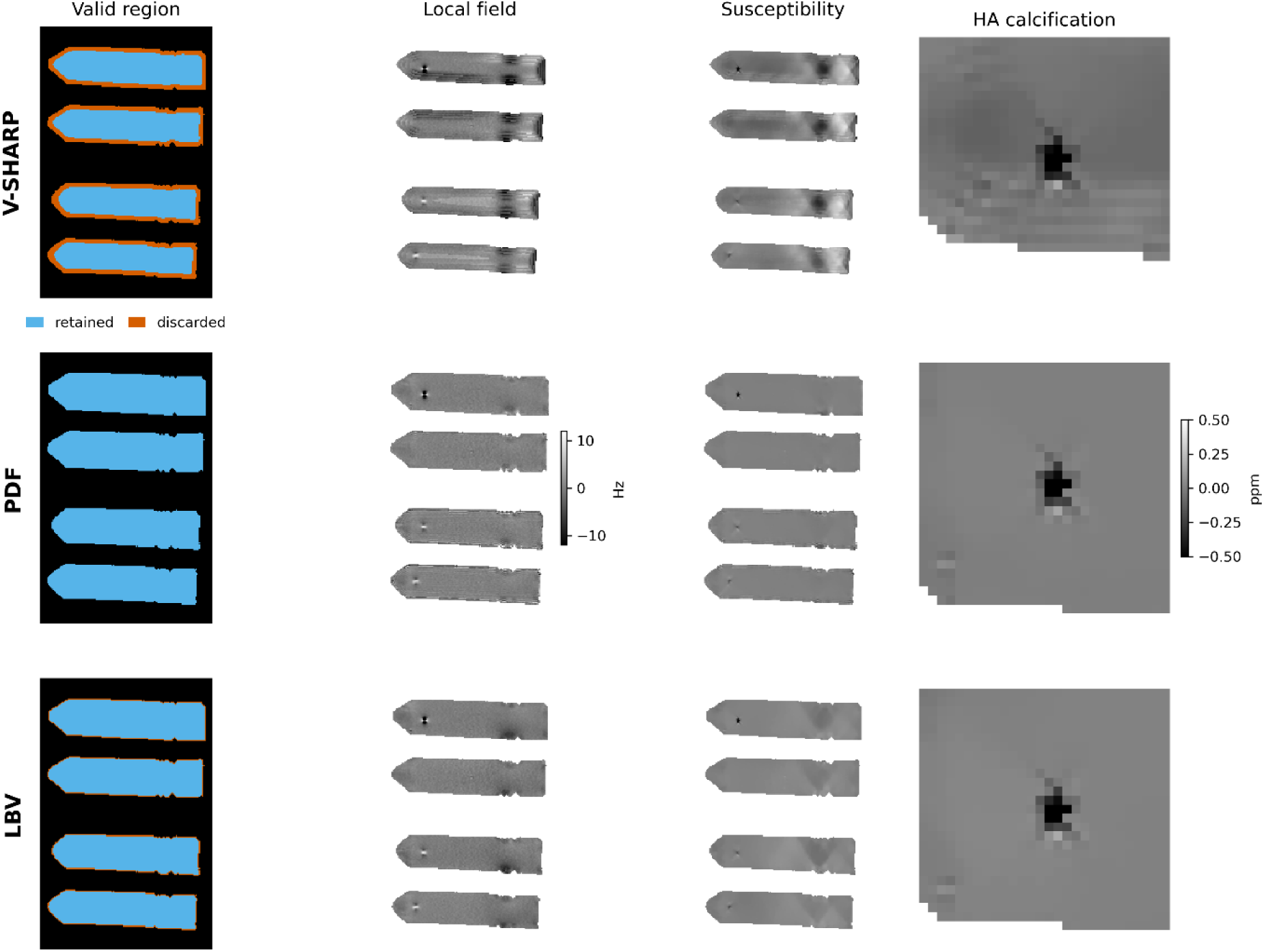
Comparison of background field removal methods on the same field map, with identical dipole inversion. Rows: V-SHARP, PDF and LBV (MEDI toolbox) (J. Liu et al., 2012). Columns: the region in which each method produces valid output, the resulting local field, the reconstructed susceptibility, and a magnified view of one HA calcification. V-SHARP discards approximately 58% of the mask through spherical-mean erosion, whereas PDF retains the full region. The magnified panels show the residual banding left by V-SHARP’s multi-radius kernels around the particle, which is absent under PDF and LBV

The difference in coverage follows from what each method assumes. V-SHARP relies on the spherical mean value property, which holds only where the kernel fits entirely within the object, so a shell of at least one kernel radius is necessarily lost from the mask boundary (Özbay et al., 2017). LBV solves a boundary value problem using the field at the mask edge as its boundary condition and retains most of the volume but not the boundary region itself (Zhou et al., 2014). PDF models the background as arising from dipole sources outside the mask and fits their contribution, which requires no clearance from the boundary and retains the full masked volume (T. Liu, Khalidov, De Rochefort, et al., 2011).

Under PDF the residual local field in the measured data was uniform with distance from the tube wall, at 0.84, 0.68, 0.73 and 1.00 Hz in successive 2 mm bands from 2 to 10 mm, and the banding present in the V-SHARP maps, arising from the boundaries between regions processed at different kernel radii, is absent (**Fig. 2**). Streaking is present at the adipose-matched particle locations and is described in **Section 3.3**.

Detection outcomes were also compared under morphology-enabled dipole inversion (MEDI) (J. Liu et al., 2012), which differs from the inversion used here (**Supplementary Table S4).** In pure alginate at 0.70 mm the two agreed, with HA detected in four of four tubes and CaOx in none, and the focal dip positive at every CaOx location under both. In the relaxometry-matched formulations they diverged, for reasons not established here.

### 3.3 Detection: HA and CaOx across gels and resolutions

HA met the detection criterion in 19 of 24 tube measurements and CaOx in four, across all gel formulations and resolutions (**Table 5**, **Fig. 5**).

**Fig. 5.**
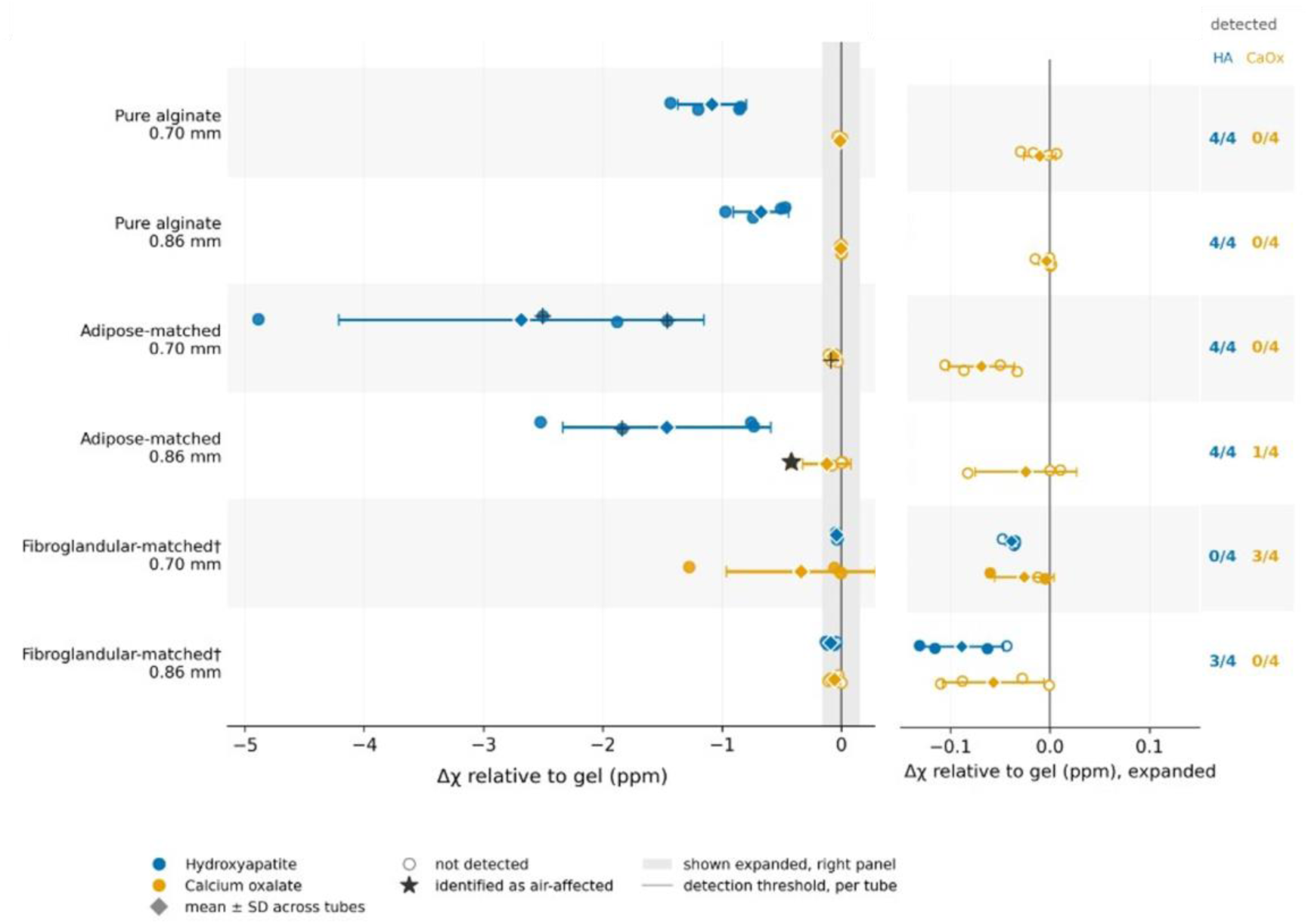
Susceptibility relative to the gel reference for HA and CaOx, by formulation and resolution. Circles are individual tubes; diamonds are condition means with the standard deviation across tubes. Open markers were not detected. Negative values are diamagnetic. The right-hand panel shows the region within ±0.15 ppm on an expanded scale, shaded on the left; short grey ticks are the per-tube detection thresholds. † Eleven of sixteen particle locations in the FGT-matched phantoms were identified as air-affected. ‡ Four locations in the adipose-matched phantoms were identified as air-affected: HA in tubes 1 and 4 at 0.70 mm and tube 2 at 0.86 mm, and CaOx in tube 2 at 0.70 mm (**Section 3.5.1**).

**Table 5.** Detection outcomes and measured susceptibility by formulation and resolution, n = 4 tubes per condition. Δχ is reported relative to the gel reference in the same tube, as mean ± standard deviation across tubes. Δχ_peak_, the 5th percentile within the particle region, is the primary metric and the quantity to which the detection criterion is applied; Δχ_mean_ is reported alongside. Cohen’s d compares HA against CaOx on Δχpeak and is descriptive at n = 4 rather than inferential. Detection requires both that |Δχ_peak_| exceeds three times the gel reference standard deviation and that a focal diamagnetic minimum is present (**Section 2.8.3**); individual measurements, with both criteria shown separately, are given in **Supplementary Table S6**.

| Formulation | Resolution | HA det. | CaOx det. | $\Delta\chi_{\text{peak}}$ HA (ppm) | $\Delta\chi_{\text{peak}}$ CaOx (ppm) | $\Delta\chi_{\text{mean}}$ HA (ppm) | $\Delta\chi_{\text{mean}}$ CaOx (ppm) | Cohen's d |
| --- | --- | --- | --- | --- | --- | --- | --- | --- |
| Pure alginate | 0.70 mm | 4/4 | 0/4 | $-1.084 \pm 0.286$ | $-0.010 \pm 0.016$ | $-0.262 \pm 0.064$ | $+0.022 \pm 0.013$ | -5.31 |
| Pure alginate | 0.86 mm | 4/4 | 0/4 | $-0.674 \pm 0.233$ | $-0.003 \pm 0.008$ | $-0.142 \pm 0.016$ | $+0.005 \pm 0.001$ | -4.06 |
| Adipose-matched <sup>a</sup> | 0.70 mm | 4/4 | 0/4 | $-2.683 \pm 1.532$ | $-0.069 \pm 0.033$ | $+0.052 \pm 0.698$ | $+0.049 \pm 0.049$ | -2.41 |
| Adipose-matched <sup>a</sup> | 0.86 mm | 4/4 | 1/4 | $-1.464 \pm 0.874$ | $-0.123 \pm 0.202$ | $+0.028 \pm 0.384$ | $+0.055 \pm 0.090$ | -2.11 |
| FGT-matched <sup>b</sup> | 0.70 mm | 0/4 | 3/4 | $-0.038 \pm 0.006$ | $-0.338 \pm 0.625$ | $+0.112 \pm 0.052$ | $+0.127 \pm 0.154$ | +0.68<sup>c</sup> |
| FGT-matched <sup>b</sup> | 0.86 mm | 3/4 | 0/4 | $-0.088 \pm 0.042$ | $-0.057 \pm 0.051$ | $+0.033 \pm 0.021$ | $+0.084 \pm 0.107$ | -0.66 |

<sup>a</sup> Four of the sixteen particle locations in this formulation were identified as air-affected, three of them HA (**Section 3.5.1**). $\Delta\chi_{\text{mean}}$ is paramagnetic where $\Delta\chi_{\text{peak}}$ is strongly diamagnetic; this is addressed in **Section 3.5.1**.
<sup>b</sup> Eleven of the sixteen particle locations in this formulation were identified as air-affected (**Section 3.5.1**).
<sup>c</sup> Cohen's d is positive here because CaOx is more diamagnetic than HA in this condition.

In pure alginate the separation was complete, with no overlap between the two mineral types at either resolution. The gel reference standard deviation in pure alginate at 0.70 mm, over a region matched in size to the particle region, was 0.0011 ppm, giving a detection threshold of 0.0033 ppm. Detection outcomes were identical under all three background field removal methods (**Section 3.2**).

CaOx appeared as a magnitude signal void at every location, of lesser depth and extent than HA: 43 ± 6% against 27 ± 4% of gel signal at 0.70 mm, and 1.9 mm against 2.5 mm in equivalent diameter. The focal dip at every CaOx location in pure alginate at 0.70 mm was positive, from +0.005 to +0.083 ppm, and positive at three of four at 0.86 mm. No location in pure alginate was identified as air-affected (**Section 3.5.1).**

Streaking is visible in the susceptibility maps at the adipose-matched particle locations, extending several voxels from the particle in a cross pattern (**Supplementary Fig. S5**). It is not confined to the air-affected locations and is absent at the pure alginate and fibroglandular-matched locations reconstructed from the same pipeline. The adipose-matched particles are the strongest sources measured, at −2.683 ± 1.532 ppm against −1.084 ± 0.286 ppm in pure alginate (**Table 5**) and streaking from single-orientation dipole inversion scales with source strength. This accounts for the paramagnetic Δχmean at those locations while Δχpeak remains strongly diamagnetic (**Section 3.5.1**).

Three of the four CaOx detections were in the FGT-matched phantoms, in which eleven of sixteen particle locations were identified as air-affected (**Section 3.5.1**); one of the three, in tube 3 at 0.70 mm, was itself among those locations. The remaining two are not accounted for by the air criterion.

The fourth CaOx detection, in adipose-matched tube 3 at 0.86 mm, gave Δχpeak of −0.419 ppm with a diamagnetic focal dip of −0.376 ppm, and is also not accounted for by the air criterion. The adjacent HA particle is not the cause: it lies 20 mm away, and the tube with the closest pairing in this formulation, 15 mm with HA at −4.89 ppm, gave a CaOx value of −0.087 ppm. The origin of these three measurements could not be determined.

#### 3.3.1 Dependence on gel formulation

Detection was complete in pure alginate at both resolutions. In the adipose-matched phantoms HA was detected in all eight measurements, but three of the eight HA locations were subsequently identified as air-affected (**Section 3.5.1**), so the detection there cannot be attributed to the mineral alone. Detection was incomplete in the FGT-matched formulation, at three of eight, and was lower at the finer resolution (0 of 4 at 0.70 mm against 3 of 4 at 0.86 mm). Measured HA susceptibility differed by a factor of 28 between pure alginate and the FGT-matched phantoms (−1.084 against −0.038 ppm at 0.70 mm).

Eleven of the sixteen particle locations in the FGT-matched phantoms were identified as air-affected (**Section 3.5.1**).

#### 3.3.2 Dependence on resolution

Measured susceptibility fell by a factor of 1.6 in pure alginate and 1.8 in the adipose-matched gel between 0.70 and 0.86 mm isotropic resolution (**Table 5**), against a ratio of voxel volumes of 1.85. The FGT-matched formulation is excluded here, since eleven of its sixteen particle locations were air-affected (**Section 3.5.1**) and the measured susceptibility does not reflect the mineral. The adipose-matched ratio includes three HA locations identified as air-affected.

The digital twin gives a comparable reduction. Recovery measured as Δχ_peak_ fell from 82.5% to 57.7% of the input susceptibility between the two resolutions, a ratio of 1.55. Measured as Δχ_mean_ the twin ratio is 2.03, reflecting the greater dilution of the region average at coarser resolution.

#### 3.3.3 Bounding the CaOx susceptibility

Detection sensitivity was established by simulation. Phantoms were constructed across CaOx susceptibilities from −1.82 to −0.02 ppm, with HA held at −2.0 ppm as an internal control, and the detection criterion of **Section 2.8.3** applied to each (**Supplementary Table S7**). Detection was complete down to −0.10 ppm, partial at −0.05 ppm (2 of 4 tubes) and absent at −0.02 ppm. HA was detected in all four tubes in every condition.

Since CaOx was not detected in any measurement in pure alginate, its susceptibility relative to the gel is smaller in magnitude than approximately 0.05 ppm.

#### 3.3.4 Dependence on particle size

The size at which detection fails was established by simulation across six equivalent diameters from 0.3 to 2.0 mm, with HA held at −2.0 ppm (**Section 2.6.2**, **Fig. 6**). Each point comprises 20 particles from five independent realisations. The 50% detection diameter was 0.38 mm (95% CI 0.35 to 0.43) at 0.70 mm isotropic resolution, 0.66 mm (0.59 to 0.75) at 1.00 mm, and 1.07 mm (0.94 to 1.19) at 1.50 mm. The two coarser resolutions were simulated rather than acquired, as representative of what a clinical breast protocol would achieve. Repeating the sweep with signal-to-noise held constant across resolutions moved the 1.50 mm floor by 0.12 mm, so the size dependence is governed by PVE rather than noise.

**Fig. 6.**
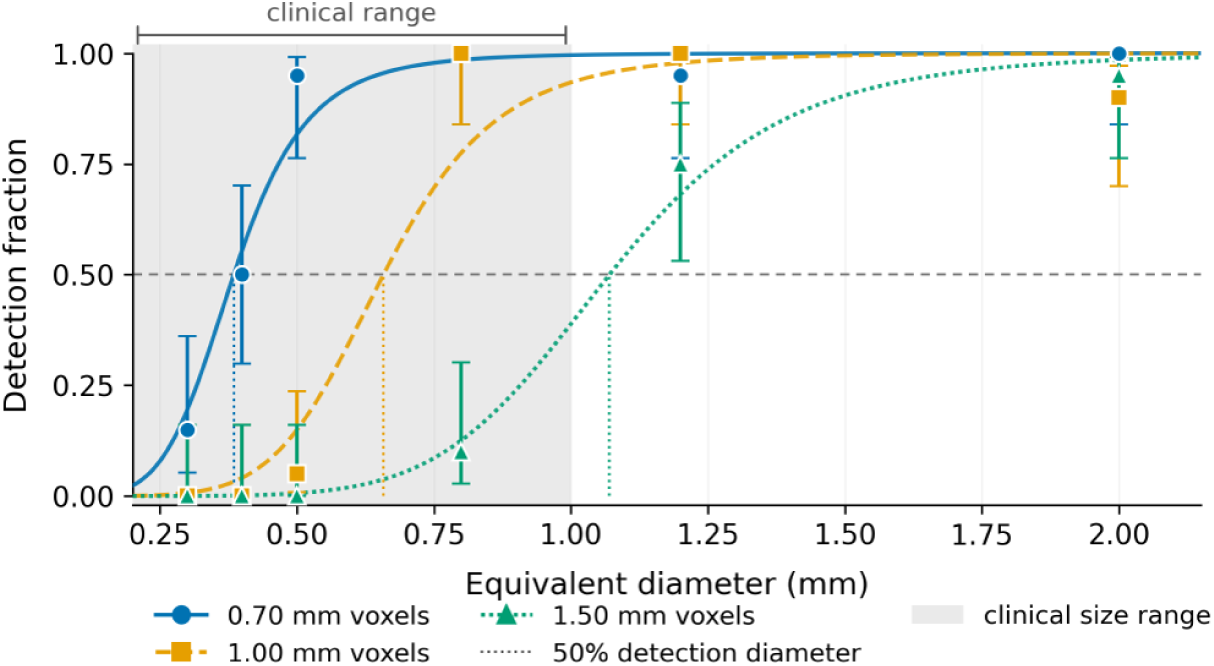
Detection of simulated HA particles against particle size at three voxel sizes. Spheroidal particles of susceptibility −2.0 ppm relative to gel were placed in the digital twin phantom and processed with the pipeline used for the measured data (**Section 2.6.2**). Each point is 20 particles, from five independent realisations of the four modelled tubes. Curves are binomial maximum-likelihood logistic fits in log diameter; dotted lines mark the 50% detection diameter. Error bars are 95% Wilson intervals, asymmetric by construction near 0 and 1; non-monotonicity between adjacent points is consistent with binomial variation at n = 20. Shading marks the clinical microcalcification range, 0.1 to 1.0 mm. Only the 0.70 mm resolution was acquired experimentally.

Applied at 1000 particle-free gel locations in the same datasets, with identical ROI geometry and gel reference, the detection criterion produced a single positive, a false positive rate of 0.1% (95% CI 0.0 to 0.6%).

The simulated particles are solid, whereas the granules used experimentally are porous (**Section 4.1**), so these diameters are a lower bound on the size required for detection.

### 3.4 R2*

R2* was elevated at both mineral types relative to the gel reference in the same tube (**Fig. 7**, **Table 6**). In pure alginate, elevation was greater for HA than for CaOx by a factor of 4.5 at 0.70 mm and 6.6 at 0.86 mm. The separation was smaller in the doped formulations, falling to 2.4 in adipose-matched and 1.3 in FGT-matched gel at 0.70 mm (**Section 4.4**).

**Fig. 7.**
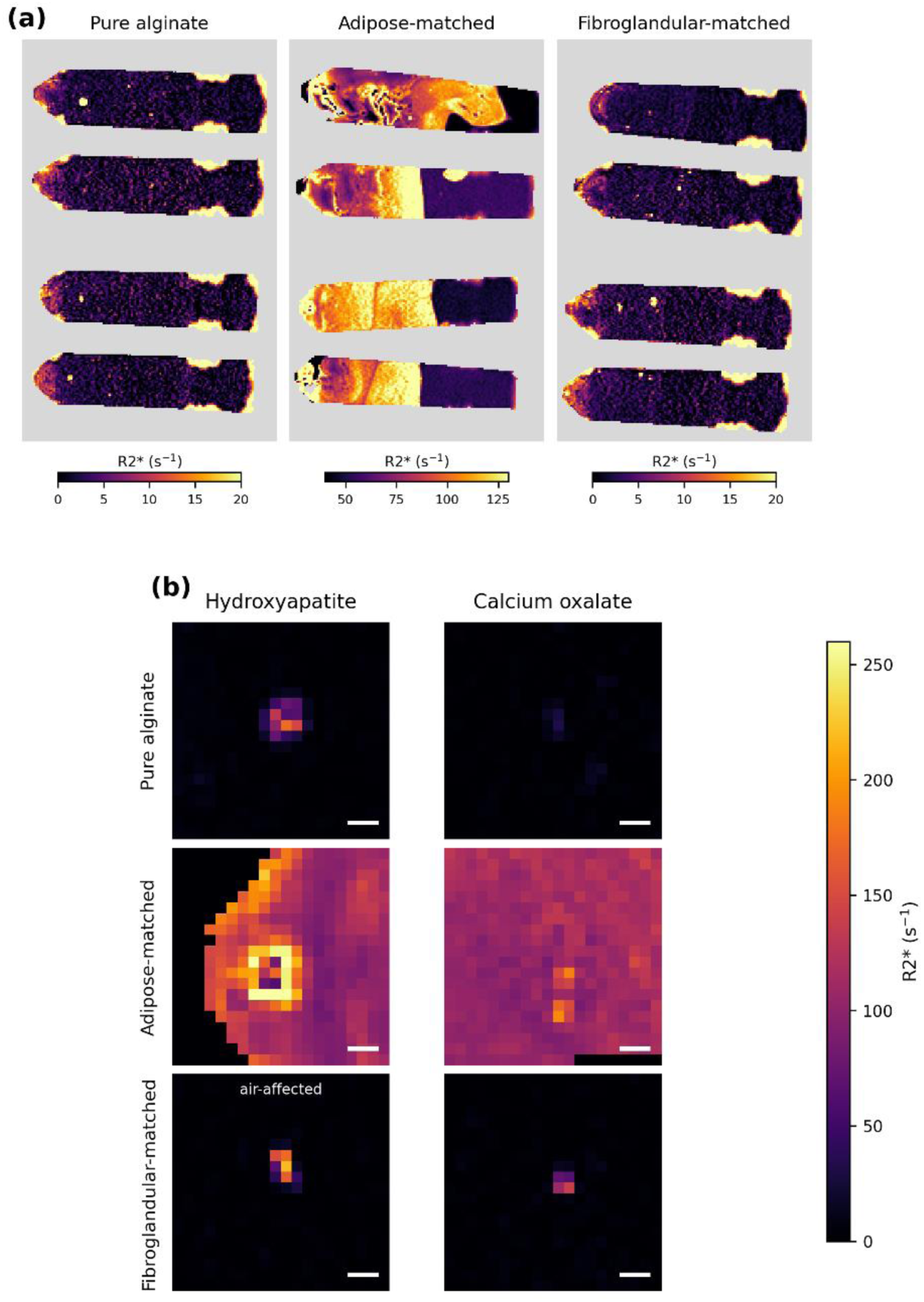
R2* in the measured phantoms. (a) One slice through each of the twelve tubes, grouped by formulation, each windowed separately because the adipose-matched gel baseline is an order of magnitude above the others; particle voxels exceed the displayed maximum and appear as small bright regions. Vertical boundaries in the adipose-matched tubes correspond to the partial-gelation embedding layers (**Section 2.3**). (b) Magnified HA and CaOx particles at 0.70 mm, from the 3rd tube of each group, chosen as closest to the condition mean of Table 6 and free of air where possible. Grey indicates voxels outside the mask. Scale bars 2 mm. All 24 particle locations at both resolutions are given in **Supplementary Figure S8**.

**Table 6.**
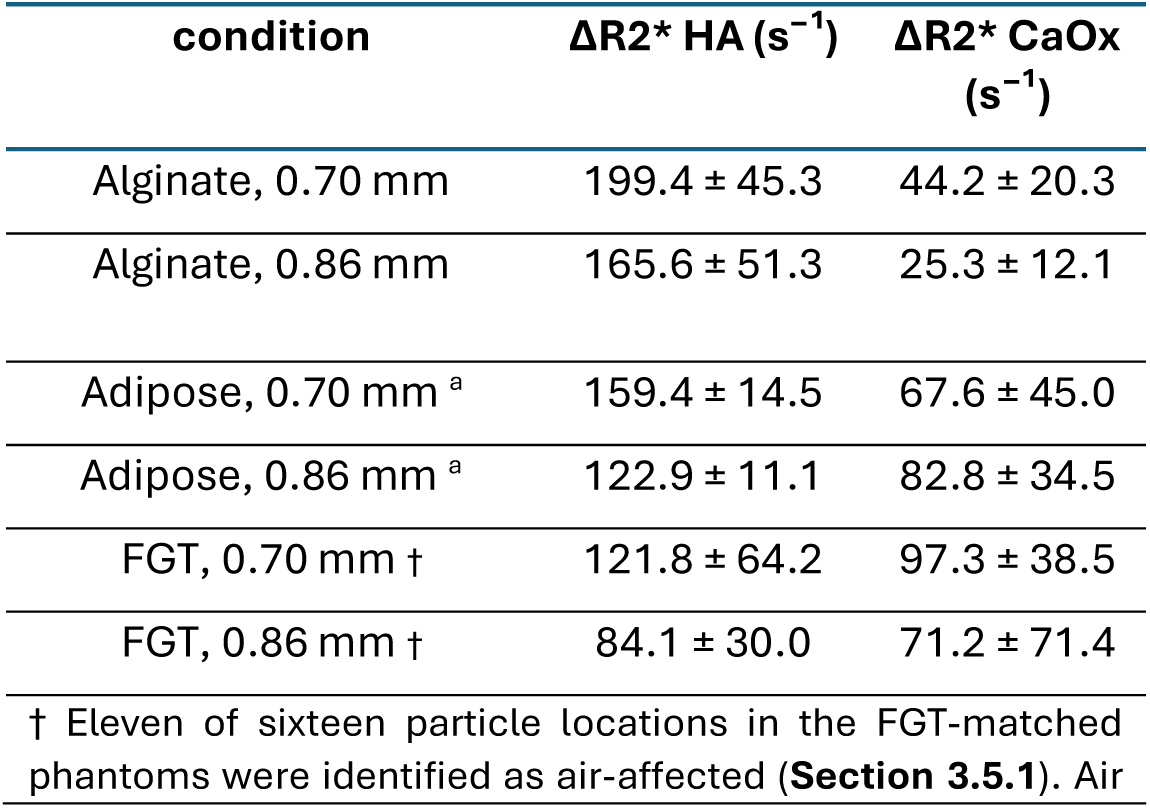
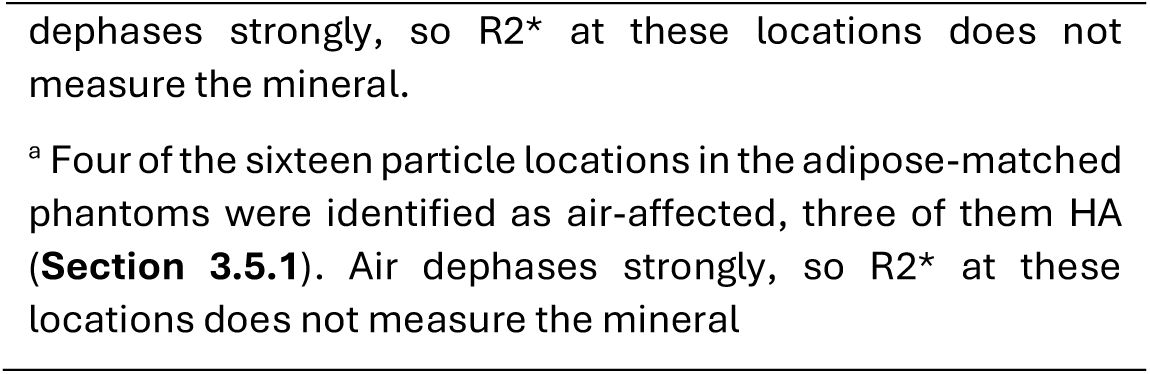
R2* elevation relative to the gel reference in the same tube, by gel formulation and resolution. Values are mean ± standard deviation across four tubes per condition.

| condition | $\Delta R2^* \text{ HA (s}^{-1}\text{)}$ | $\Delta R2^* \text{ CaOx (s}^{-1}\text{)}$ |
| --- | --- | --- |
| Alginate, 0.70 mm | $199.4 \pm 45.3$ | $44.2 \pm 20.3$ |
| Alginate, 0.86 mm | $165.6 \pm 51.3$ | $25.3 \pm 12.1$ |
| Adipose, 0.70 mm <sup>a</sup> | $159.4 \pm 14.5$ | $67.6 \pm 45.0$ |
| Adipose, 0.86 mm <sup>a</sup> | $122.9 \pm 11.1$ | $82.8 \pm 34.5$ |
| FGT, 0.70 mm † | $121.8 \pm 64.2$ | $97.3 \pm 38.5$ |
| FGT, 0.86 mm † | $84.1 \pm 30.0$ | $71.2 \pm 71.4$ |

† Eleven of sixteen particle locations in the FGT-matched phantoms were identified as air-affected (**Section 3.5.1**). Air
dephases strongly, so $R2^*$ at these locations does not measure the mineral.
<sup>a</sup> Four of the sixteen particle locations in the adipose-matched phantoms were identified as air-affected, three of them HA (**Section 3.5.1**). Air dephases strongly, so $R2^*$ at these locations does not measure the mineral

The truncation criterion of **Section 2.7** excluded no echoes at these signal levels: the median number used was seven of seven in every dataset, and fitted rates were unchanged when all echoes were included unconditionally.

### 3.5 Confounding factors

#### 3.5.1 Trapped air

Fifteen of the 48 particle locations, 31%, were identified as air-affected: none in pure alginate, four of sixteen in the adipose-matched phantoms, and eleven of sixteen in the FGT-matched. The identifications differ in how far they exceed their threshold: the adipose-matched locations by 0.5 to 58 Hz, the FGT by 0.1 to 7.0 Hz. Measured against the matched null of **Section 2.9**, whose standard deviation is 0.23 Hz, the FGT identifications lie between 8 and 37 standard deviations from zero and the adipose ones between 80 and 264. Shifting the threshold by ±0.5 Hz changes the total from 13 to 18, with the FGT contribution moving from nine to thirteen; no location in pure alginate is identified at any threshold in that range.

Magnitude signal voids at the affected FGT locations were of similar depth to those in pure alginate, 35% against 27% of gel signal, so the particles are present at those positions.

Three of the eight adipose-matched HA locations were identified as air-affected, at +18.6, +27.6 and +60.7 Hz, and those three are the locations at which the region mean is paramagnetic. The focal dips there remain strongly diamagnetic, from −0.31 to −3.06 ppm, so the particle lies at the centre of the region while the surrounding field carries a large paramagnetic contribution from the adjacent air.

The effect of an adjacent air inclusion on recovered susceptibility was examined in the digital twin. Inclusions of 2.5 mm diameter were placed at varying separations from an HA particle, with one tube per condition left without an inclusion as a control (**Table 7**). An adjacent inclusion inverted the sign of the recovered susceptibility in the FGT-matched formulation, returning +0.009 ppm against a partial-volume ground truth of −0.249 ppm, but not in pure alginate, where recovery was 71.9%. With one tube per condition these are single measurements: sign inversion was observed in one of the two, and the conditions under which it occurs are not established. Gel reference variability rose 6 to 14-fold in tubes containing inclusions and 2-fold in a tube without one in the same phantom, so the presence of an inclusion disturbs the reconstruction across the tube rather than only at the particle.

**Table 7.** Recovered HA susceptibility in the digital twin as a function of separation from a 2.5 mm air inclusion, as recovery against partial-volume ground truth. One tube per condition, so differences between adjacent separations are single measurements and are not interpretable as a trend.

| separation | alginate | FGT-matched |
| --- | --- | --- |
| none (control) | 95.7% | 98.1% |
| 4 mm | 95.0% | 86.7% |
| 2 mm | 89.3% | 98.4% |
| adjacent | 71.9% | -3.6% |

#### 3.5.2 Receive sensitivity

The receive sensitivity profile, measured from the gel, fell to between 0.61 and 0.69 at the edge of the phantom relative to the centre depending on formulation (**Section 2.6.1**). Recovered susceptibility changed by less than 1% across this gradient in the digital twin.

#### 3.5.3 Particle position

Exchanging the HA and CaOx positions in the digital twin, with all other parameters held fixed, gave recovery against partial-volume ground truth of 94.0% for HA in both configurations, and 94.3% and 94.5% for CaOx. The asymmetry between the two minerals therefore does not depend on placement within the tube. These values exceed the 89.8% of **Table 3** because they derive from a single particle and noise realisation rather than the 80 measurements averaged there.

## 4. Discussion

HA was detected by QSM in every measurement in pure alginate and CaOx in none, at both resolutions. The distinction is categorical rather than quantitative: the CaOx granules produced no measurable susceptibility contrast, so susceptibility mapping alone separates HA from its absence rather than the two minerals from one another. R2* was elevated at both particle types and distinguished them by magnitude, so the two contrasts together resolve three cases from a single multi-echo acquisition: HA, CaOx, and no particle.

### 4.1 The CaOx result is an absence of contrast, not a failure to detect

CaOx was not detected in any measurement in pure alginate. Four observations indicate that this reflects an absence of susceptibility contrast rather than a source too weak to resolve.

First, the detection threshold in pure alginate at 0.70 mm was 0.0033 ppm, some three hundred times smaller than the −1.08 ± 0.13 ppm reported for CaOx in renal calculi (Mücke et al., 2026), and simulation places the limit at approximately 0.05 ppm relative to the gel, still twenty times below it. A source of the reported magnitude would have been detected comfortably.

Second, the CaOx used here differs from that in renal calculi. The reagent is the dihydrate, as in breast Type I calcifications (Haka et al., 2002), whereas renal calculi are predominantly the monohydrate and often contain apatite (Radvar et al., 2021). The particles are also compression granules rather than dense crystalline stones; taking the solid fraction of 0.77 reported for this protocol (Ghammraoui et al., 2021), the bound on the solid mineral is an estimated 0.065 ppm, still seventeen times smaller than the renal calculi value. Whether the difference reflects hydration state, composition or granule structure is not established here, and the susceptibility of this reagent was not measured independently (**Section 4.8**).

Third, the focal dip was positive at every CaOx location in pure alginate at 0.70 mm, indicating a centre voxel more paramagnetic than its surroundings, and positive at three of four at 0.86 mm. None of these locations was identified as air-affected (**Section 3.5.1**).

Lastly, if the granule pore volume were air-filled, HA at 40% porosity would read +3.0 ppm and CaOx at 23% would read +1.3 ppm, both strongly paramagnetic. HA is measured at −1.084 ppm at 0.70 mm, so the pores are predominantly wetted. On that basis the residual paramagnetic signal at CaOx sites, +0.062 ± 0.021 ppm at 0.70 mm, corresponds to approximately 0.6% of the granule volume as air, or about 3% of its pore space. Since +0.062 ppm is a reconstructed, partial-volume diluted value, this is a lower bound on the air fraction.

Together these indicate granules whose pores are predominantly wetted and whose mineral susceptibility lies below the detection limit, rather than a source present in the phantom but not recovered by the measurement.

### 4.2 Convergence with an independent pipeline

The reconstruction used here converges closely with that of the renal calculus study against which the CaOx result is compared (Mücke et al., 2026), despite being implemented independently. Both mask by thresholding the magnitude and eroding by two to three voxels, unwrap with ROMEO, remove background fields by projection onto dipole fields, and reference susceptibility to the surrounding medium. Both select the dipole inversion regularisation weight by optimisation rather than adopting a default, and both validate the reconstruction against a forward-simulated numerical phantom. The remaining difference is the inversion algorithm itself, morphology-enabled dipole inversion there and total variation regularised ADMM here. Detection outcomes under the two agreed in pure alginate (**Section 3.2**).

### 4.3 R2* provides confirmation independent of the susceptibility pathway

R2* is derived from the decay of the multi-echo magnitude and involves no phase unwrapping, no background field removal and no dipole inversion. It is therefore unaffected by every stage at which the susceptibility measurement could fail, so the two contrasts agree without sharing any processing step.

In pure alginate, R2* separated the two mineral types by a factor of 4.5 at 0.70 mm and 6.6 at 0.86 mm, in the same direction as the susceptibility measurement. The separation is smaller in the doped formulations, for the reasons given in **Section 4.4**.

The same asymmetry has been measured a third way. Blooming artefact extent at 9.4 T has been reported as more than twice as large for HA as for CaOx (Mustafi et al., 2015). Blooming arises from intravoxel dephasing, the same mechanism that elevates R2*. Susceptibility, R2* and blooming extent, measured at different field strengths and in different laboratories, all find HA producing the stronger effect.

The two contrasts have different sensitivities. Susceptibility mapping detected HA and not CaOx, while R2* was elevated for both. Taken together they resolve three cases rather than two in pure alginate: a focal susceptibility source indicates HA, elevated R2* without one indicates CaOx, and neither indicates that no particle is present. The mechanism by which CaOx elevates R2* is not established here: its susceptibility is too small to account for the elevation by bulk field perturbation alone, so the granule structure itself, rather than the mineral, is the likely source.

Both are obtained from the same multi-echo acquisition, R2* from the magnitude and susceptibility from the phase, so the second contrast costs no additional scan time.

### 4.4 Differences between formulations

Measured HA susceptibility differed by a factor of 28 between the pure alginate and FGT-matched phantoms (**Section 3.3.1**). The reconstruction cannot account for this: it recovers the same fraction of a known input in every formulation, 18.2%, 18.1% and 18.2% respectively (**Section 3.1**), despite simulated baseline R2* spanning 5.9 to 84.2 s⁻¹ and SNR spanning 27.3 to 44.0.

The difference therefore originates in the phantoms. Eleven of sixteen FGT-matched particle locations were identified as air-affected, all lying 8 to 37 standard deviations from the matched null (**Section 3.5.1**), and an adjacent inclusion inverted the sign of the recovered susceptibility in the digital twin, though with one tube per condition that observation is a single measurement (**Section 3.5.1**).

The R2* separation ratio fell from 4.5 in pure alginate to 2.4 in adipose-matched and 1.3 in FGT-matched gel at 0.70 mm. The reduction has a different cause in each. In the FGT-matched phantoms it is trapped air, which dephases strongly and elevates R2* at affected locations regardless of the mineral present.

In the adipose-matched phantoms the reduction has two contributions. Three of the eight HA locations were identified as air-affected (**Section 3.5.1**), so R2* there carries a contribution that is not the mineral. At the remaining locations the baseline itself is the limit: measured gel R2* ranges from 64 to 103 s⁻¹ between tubes against 2.3 s⁻¹ measured in pure alginate, so the particle contribution sits on a larger and much more variable background, and the gel reference standard deviation over the whole gel is correspondingly higher, 0.0275 against 0.0056 ppm in susceptibility. The adipose-matched gel also varies within tubes, at boundaries corresponding to the embedding layers (**Fig. 7**), so the dopant is incompletely mixed between layers.

### 4.5 Air retained within the granules

Granules produced by compression granulation retain substantial pore volume, approximately 40% for HA and 23% for CaOx at the compression load used (Ghammraoui et al., 2021). Whether those pores fill with the surrounding medium or retain air determines the sign of the apparent susceptibility: a granule with wetted pores reads as a diluted mineral, while one retaining air reads paramagnetic, since air is approximately +9.4 ppm relative to water, against −1.25 to −2.0 ppm for HA and below 0.065 ppm for CaOx.

Gels were centrifuged before assembly, which removes entrained air from the gel but cannot displace air from within the granules, nor prevent its entrapment when a dry granule is placed onto a gelled surface and covered.

Dopant concentration does not account for the pattern. The adipose-matched gel carries the higher divalent cation load, at 4.23 mM against 1.56 mM, yet showed four affected locations against eleven in the FGT-matched. The adipose identifications are the larger ones, exceeding their threshold by 0.5 to 58 Hz against 0.1 to 7 Hz in the FGT, so neither count nor magnitude follows the dopant concentration.

Eleven of sixteen particle locations in the FGT-matched phantoms were affected, despite identical crosslinking chemistry and gel properties comparable to pure alginate: baseline R2* of 2.1 s⁻¹ against 2.3 s⁻¹, whole-gel susceptibility variability of 0.0044 ppm against 0.0056 ppm, and magnitude coefficient of variation of 9.8% against 7.9%.

Affected measurements were identified by the sign of the local field along B₀, measured before dipole inversion (**Section 2.9**). The focal dip in the susceptibility map shows the same pattern, but that quantity depends on the inversion: under morphology-enabled dipole inversion the pure alginate CaOx dips rise from +0.062 ± 0.021 to +0.156 ± 0.060 ppm (**Supplementary Table S3**), a factor of 2.5. Any threshold placed on that quantity would therefore be specific to one reconstruction. The local field criterion is unaffected by that choice.

Related observations have been reported. A phantom study of renal calculi reported a weakly paramagnetic shell at the agar–stone boundary and excluded those regions from its volumes of interest (Mücke et al., 2026); air cavities have been observed around HA samples in gelatine (Kan et al., 2022); and air-free fabrication of calcium mineral phantoms has been described (Mustafi et al., 2015).

Air was identified at 15 of 48 particle locations, 31% of the measurements, and at none in pure alginate. This is a finding about compression granulation rather than about these phantoms specifically. Granules with 23 to 40% pore volume can retain air when placed dry onto a gelled surface, though this occurred in the doped formulations only and the reason for that difference is not established.

### 4.6 Resolution dependence

Measured susceptibility fell by a factor of 1.6 between 0.70 and 0.86 mm (**Section 3.3.2**), against a ratio of voxel volumes of 1.85 and a simulated reduction of 1.55. The agreement, and the twin’s attribution of the change to voxel sampling rather than to the reconstruction, indicates that the resolution dependence is geometric.

Susceptibility values for particles at this scale are therefore resolution-dependent and should be reported with the acquisition voxel size.

The same dependence sets the size at which detection fails. Simulation places the 50% detection diameter at 0.38 mm at 0.70 mm resolution, rising to 0.66 mm at 1.00 mm and 1.07 mm at 1.50 mm (**Section 3.3.4**). Clinical microcalcifications span 0.1 to 1.0 mm, so a 1.50 mm acquisition reaches almost none of that range while a 0.70 mm acquisition reaches most of it. Since the simulated particles are solid and granules used in this study are porous, these are a lower bound on the size required.

### 4.7 Confounds tested and found not limiting

Receive sensitivity varies by a factor of 1.6 across the phantom, and the head coil used is not the coil that would be used clinically. Neither affects the measurement materially: recovered susceptibility changed by less than 1% across the measured gradient. The coil affects SNR and coverage, not the susceptibility recovered from a given signal. Particle position is likewise not a factor (**Section 3.5.3**).

### 4.8 Limitations

The digital twin models solid ellipsoidal particles. The particles used are compression granules produced from powder with a binder, and gel infiltrates the pore space, so their effective susceptibility is diluted relative to the solid mineral. The recovery figures therefore describe the voxel-sampling component of PVE and not internal dilution and are a lower bound on the total difference between mineral susceptibility and reported value. Ellipsoids are also smoother than crushed granules, and particle geometry is the dominant source of variability in the recovery measurement.

Single-orientation dipole inversion produces streaking around strong compact sources, and this is visible at the adipose-matched particle locations (**Section 3.3**). Detection outcomes were unchanged across three background field removal methods and two dipole inversions (**Section 3.2**), so the streaking does not determine which particles are detected, but it does affect the susceptibility values at those locations

The background field in the twin is a linear shim gradient of ±300 Hz. The background measured experimentally has a standard deviation of 454 Hz with approximately 30% of its variance in components that are not linear. The twin therefore presents background field removal with an easier problem than the acquisition does. This does not apply to the method comparison of **Section 3.2**, for which a third-order polynomial fit to the measured background was used in place of the synthetic shim.

Gradient nonlinearity and eddy currents are not modelled. The phantom sits within approximately 90 mm of isocentre, where gradient distortion is sub-millimetre against a 2 mm particle in 0.70 mm voxels, and the readout is monopolar, which largely avoids odd-even echo inconsistency.

Signal is synthesised with constant proton density rather than from the spoiled gradient-echo steady state, with noise scaled to the measured per-formulation signal-to-noise ratio. The steady-state signal differs by up to a factor of 2.15 between formulations owing to their differing T₁, but susceptibility is referenced to gel within the same tube, so a constant scale factor cancels. Residual transverse coherence from imperfect RF spoiling scales as sin²(α/2) = 0.017 at the flip angle used and affects gel and particle equally.

The gels contain no lipid. The formulations match breast tissue relaxation times but not composition, so the chemical shift effects present in vivo are absent (**Section 4.9**). The adipose-matched formulation therefore isolates the effect of relaxation and noise on detection rather than modelling adipose tissue for susceptibility mapping, for which the chemical shift is the dominant consideration.

The susceptibility of the CaOx reagent was not measured independently. The bound in **Section 3.3.3** rests on non-detection under a criterion validated by simulation, and an independent bulk measurement would be needed to reconcile it with the value reported for renal calculi (Mücke et al., 2026).

### 4.9 Clinical implications and future work

PVE, not susceptibility contrast, sets the limit. At 0.70 mm isotropic, a particle of the size studied here occupies approximately 12 of the 33 voxels in the measurement region, and recovery of the input susceptibility is 17%. At 0.86 mm it falls to 9%. Clinical breast MRI is acquired at coarser resolution than the phantom data here, with thicker slices, and microcalcifications are smaller than the particles used. **Section 4.6** quantifies the consequence: the 50% detection diameter rises from 0.38 mm at 0.70 mm resolution to 1.07 mm at 1.50 mm. The susceptibility values reported here should therefore be read as an upper bound on what would be recovered in vivo at clinically achievable resolutions.

Fat is the principal obstacle to applying the two-parameter scheme in vivo. Both contrasts are affected. Susceptibility mapping requires a field estimate, and fat resonates approximately 3.4 ppm from water, so a single-species fit misattributes chemical shift as off-resonance. R2* cannot be estimated by fitting raw magnitude where fat and water coexist within a voxel, since their interference modulates the signal; a chemical shift-encoded model is required. Chemical shift-encoded reconstruction addresses both (Dimov et al., 2015; Stelter et al., 2022), and has been shown to reduce streaking and increase recovered susceptibility values in a phantom containing peanut oil (Graf et al., 2025).

The fabrication finding has implications beyond this study. Compression granulation (Ghammraoui et al., 2021) is an established method for producing synthetic calcifications, and a granule retaining air can read paramagnetic, which is indistinguishable from a calcification on magnitude alone. The local field sign check described in **Section 2.9** is recommended as a routine screen for phantoms of this type.

## 5. Conclusion

HA and CaOx were distinguished in tissue-mimicking phantoms at 3 T, with complete separation in undoped alginate. The distinction is categorical rather than quantitative. The CaOx granules produced no measurable susceptibility contrast, below 0.05 ppm relative to the gel and an estimated 0.065 ppm on the solid mineral, while R2* was elevated at both particle types and separated them by a factor of 4.5 at 0.70 mm. The two contrasts, from a single acquisition, resolve three cases where either alone resolves two.

PVE rather than susceptibility contrast sets the limit: 18.2% of the input susceptibility was recovered, of which the larger loss was geometric, and the 50% detection diameter rises from 0.38 mm at 0.70 mm resolution to 1.07 mm at 1.50 mm. Air retained within the porous granules affected 31% of particle locations, a fabrication effect detectable from the sign of the local field.

## Author contributions

KM designed the study, fabricated the phantoms, implemented the reconstruction pipeline and digital twin, performed the analysis and wrote the manuscript. CAC, MTC and SWS supervised. EDV advised on MRI acquisition. All authors reviewed and approved the final manuscript.

