## Supplemental Information for "Quantitative Susceptibility Mapping for Differentiating Hydroxyapatite and Calcium Oxalate Breast Calcifications at 3T: A Phantom Study"

### S1. Regularisation weight selection

**Table S1**. Selection of the regularisation weight against known susceptibility in the digital twin phantom. RMSE values are root-mean-square error against ground truth; recovery is the mean recovered susceptibility as a percentage of the known input.

| λ | RMSE, whole tube (ppm) | RMSE, particle (ppm) | mean recovery (%) | gel SD (ppm) |
|---|---|---|---|---|
| $5 \times 10^{-5}$ | 0.0290 | 0.4339 | 91.5 | 0.0285 |
| $5 \times 10^{-4}$ | 0.0130 | 0.4011 | 94.0 | 0.0121 |
| $2 \times 10^{-3}$ | 0.0060 | 0.3934 | 91.9 | 0.0038 |
| $5 \times 10^{-3}$ | 0.0064 | 0.4593 | 85.8 | 0.0027 |

### S2. Verification of the twin forward model

The forward model was verified against cases with known analytic solutions.

**Table S2**. Verification of the digital twin forward model against analytic solutions and internal consistency requirements. Susceptibility values are relative to the surrounding medium; the cylinder is simulated at an aspect ratio of 8:1 rather than infinite.

| test | expected | computed |
|---|---|---|
| uniform sphere, interior field | 0 | 0 to numerical precision |
| sphere exterior, $r = 2a$, along and perpendicular to $B_0$ | dipole law | agrees to 6% |
| infinite cylinder, ratio of parallel to perpendicular interior field | −2 | −2.0000 |
| cylinder interior, individual values | $\pm\chi/3$, $\mp\chi/6$ | agrees to 10% |
| linearity: field ratio for doubled susceptibility | 2 | 2.000 |
| superposition of separated sources | exact | $4 \times 10^{-16}$ |
| R2* recovered from synthesized signal, 7.5 / 84.2 / 175.5 $s^{-1}$ | as specified | exact |
| ROI extraction, simulation against experimental | identical | 33 voxels |

The sphere provides the sharper geometric test, since its interior field must be exactly zero: the Lorentz term in the dipole kernel cancels the demagnetising factor in every direction, so an error in normalisation cannot be absorbed into a plausible non-zero value. The 10% deviation in the cylinder interior arises from the finite length of the simulated tube, an aspect ratio of 8:1 rather than infinite; the ratio between orientations is unaffected.

#### S3. Size-dependent detection by simulation

Table S3 gives the full results of the sweep summarised in **Section 3.3.4**: six equivalent diameters at three acquisition resolutions, 20 particles per condition from five independent realisations of four particles, with HA held at −2.0 ppm and a R2* increment of 175.5 $s^{-1}$.

**Table S*3*.** Detection and recovered susceptibility against particle size and voxel size. HA was simulated at −2.0 ppm relative to gel with a R2* increment of 175.5 $s^{-1}$, at six equivalent diameters and three acquisition resolutions, with 20 particles per condition from five independent realisations of four particles. Detection is given for both noise models: signal-to-noise scaling with voxel volume, as at fixed acquisition time, and held constant across resolutions. Peak Δχ is the fifth percentile over the region of interest, mean (SD) across particles. The threshold is three times the gel standard deviation. Recovery is the region mean as a percentage of the partial-volume ground truth.

| (mm) | d (mm) | d/v | Detected, fixed time | Detected, fixed SNR* | Peak Δχ (ppm) | Threshold (ppm) | Recovery (%)† |
|---|---|---|---|---|---|---|---|
| 0.70 | 0.3 | 0.43 | 3/20 | * | −0.009 (0.007) | 0.021 | |
| 0.70 | 0.4 | 0.57 | 10/20 | * | −0.022 (0.011) | 0.021 | |
| 0.70 | 0.5 | 0.71 | 19/20 | * | −0.053 (0.021) | 0.021 | 88 |
| 0.70 | 0.8 | 1.14 | 20/20 | * | −0.275 (0.114) | 0.022 | 90 |
| 0.70 | 1.2 | 1.71 | 19/20 | * | −0.830 (0.244) | 0.025 | 95 |
| 0.70 | 2.0 | 2.86 | 20/20 | * | −2.104 (0.292) | 0.031 | 84 |
| 1.00 | 0.3 | 0.30 | 0/20 | 0/20 | −0.001 (0.001) | 0.034 | |
| 1.00 | 0.4 | 0.40 | 0/20 | 0/20 | −0.004 (0.003) | 0.034 | |
| 1.00 | 0.5 | 0.50 | 1/20 | 2/20 | −0.013 (0.007) | 0.034 | |
| 1.00 | 0.8 | 0.80 | 20/20 | 20/20 | −0.082 (0.024) | 0.034 | 87 |
| 1.00 | 1.2 | 1.20 | 20/20 | 20/20 | −0.347 (0.242) | 0.036 | 87 |
| 1.00 | 2.0 | 2.00 | 18/20 | 19/20 | −1.050 (0.345) | 0.042 | 89 |
| 1.50 | 0.3 | 0.20 | 0/20 | 0/20 | −0.002 (0.004) | 0.057 | |
| 1.50 | 0.4 | 0.27 | 0/20 | 0/20 | −0.002 (0.004) | 0.057 | |
| 1.50 | 0.5 | 0.33 | 0/20 | 0/20 | −0.003 (0.004) | 0.057 | |
| 1.50 | 0.8 | 0.53 | 2/20 | 0/20 | −0.022 (0.009) | 0.057 | |
| 1.50 | 1.2 | 0.80 | 15/20 | 11/20 | −0.083 (0.019) | 0.057 | 69 |
| 1.50 | 2.0 | 1.33 | 19/20 | 18/20 | −0.378 (0.136) | 0.069 | 79 |

* At 0.70 mm the two noise models coincide by construction, signal-to-noise being referenced to that resolution.

† Recovery is reported only where the particle fills more than 10% of the region of interest. Below that the ratio divides by a near-zero ground truth and is uninformative

Expressed as a fraction of the voxel size, the 50% detection diameters are 0.55, 0.66 and 0.71 at 0.70, 1.00 and 1.50 mm respectively, so the dependence is largely geometric. It is not purely so: the ratio rises by about 30% across a twofold change in voxel size, and while the intervals for the finest and coarsest resolutions do not overlap, the intermediate resolution overlaps both. Part of the residual dependence arises because the detection threshold itself scales with voxel size, the gel standard deviation being 0.007, 0.011 and 0.019 ppm at the three resolutions. Repeating the sweep with signal-to-noise held constant across resolutions, rather than allowing it to scale with voxel volume as it would at fixed acquisition time, moved the 50% detection diameter at 1.50 mm from 1.07 to 1.19 mm and left 1.00 mm unchanged, indicating that the size dependence is governed by partial volume rather than by noise.

The criterion is highly specific. Applied at 1000 particle-free gel locations in the same datasets, using identical region geometry and gel reference, it produced a single positive, a false positive rate of 0.1% (95% CI 0.0 to 0.6%).

At the condition common to this sweep and the established digital twin, 2.0 mm particles at 0.70 mm resolution, the region mean recovered 28.7 ± 2.6% of the input susceptibility and 83.8 ± 7.5% of the partial-volume ground truth, consistent with the established implementation. Across the sweep, recovery relative to the partial-volume ground truth was 83 to 95% at 0.70 and 1.00 mm and 69 to 79% at 1.50 mm. The one departure from this pattern is at 0.70 mm, where the 2.0 mm particles recovered 84 ± 8% against 95% at 1.2 mm, consistent with the signal-void core of a particle much larger than a voxel providing no usable phase to the inversion.

Four particles at diameters of 1.2 mm and above failed the focal dip requirement despite peak values 10 to 30 times the threshold, three of them returning a positive focal dip. The requirement compares a fixed central voxel against a shell two voxels away, which is unreliable when a particle straddles voxel boundaries at diameter-to-voxel ratios near two. The five further misses at 1.2 mm and 1.50 mm resolution are of a different kind: their peak values are close to the threshold, so those particles are genuinely marginal rather than misclassified. Neither affects the detection floors or any measured result.

#### S4. Inversion Comparison

**Table S4**. Detection outcomes and susceptibility measurements under two dipole inversion methods, applied to identical local field maps. Total variation regularised ADMM is the method used throughout; morphology-enabled dipole inversion (MEDI) was applied with its regularisation weight selected against known susceptibility in the digital twin phantom, and with each tube inverted separately. Values are mean ± standard deviation across four tubes per condition. Δχpeak is the 5th percentile within the particle region minus the mean gel reference; the focal dip is the centre voxel minus the mean of voxels two voxels away.

| Formulation | Res (mm) | | TV-ADMM | MEDI |
|---|---|---|---|---|
| Pure alginate | 0.70 | HA detected | 4/4 | 4/4 |
| | | CaOx detected | 0/4 | 0/4 |
| | | HA $\Delta\chi_{peak}$ (ppm) | −1.084 ± 0.286 | −0.628 ± 0.113 |

| | | | | |
|---|---|---|---|---|
| | | CaOx focal dip (ppm) | +0.062 ± 0.021 | +0.156 ± 0.060 |
| | | gel SD (ppm) | 0.0011 | 0.0008 |
| Pure alginate | 0.86 | HA detected | 4/4 | 3/4 |
| | | CaOx detected | 0/4 | 1/4 |
| | | HA $\Delta\chi_{peak}$ (ppm) | −0.674 ± 0.233 | −0.488 ± 0.245 |
| | | CaOx focal dip (ppm) | +0.008 ± 0.008 | +0.059 ± 0.056 |
| | | gel SD (ppm) | 0.0009 | 0.0010 |
| Adipose-matched | 0.70 | HA detected | 4/4 | 0/4 |
| | | CaOx detected | 0/4 | 0/4 |
| | | HA $\Delta\chi_{peak}$ (ppm) | −2.683 ± 1.532 | −0.128 ± 0.681 |
| | | gel SD (ppm) | 0.0060 | 0.0042 |
| Adipose-matched | 0.86 | HA detected | 4/4 | 0/4 |
| | | CaOx detected | 1/4 | 0/4 |
| | | HA $\Delta\chi_{peak}$ (ppm) | −1.464 ± 0.874 | −0.012 ± 0.046 |
| | | gel SD (ppm) | 0.0057 | 0.0083 |
| FGT-matched | 0.70 | HA detected | 0/4 | 0/4 |
| | | CaOx detected | 3/4 | 1/4 |
| | | HA $\Delta\chi_{peak}$ (ppm) | −0.038 ± 0.006 | +0.002 ± 0.006 |
| | | gel SD (ppm) | 0.0023 | 0.0025 |
| FGT-matched | 0.86 | HA detected | 3/4 | 0/4 |
| | | CaOx detected | 0/4 | 0/4 |
| | | HA $\Delta\chi_{peak}$ (ppm) | −0.088 ± 0.042 | −0.010 ± 0.016 |
| | | gel SD (ppm) | 0.0013 | 0.0006 |

Detection outcomes agreed in pure alginate at 0.70 mm, the condition in which the gel reference standard deviation is lowest, and no particle location was identified as air affected. The focal dip at every CaOx location was positive under both inversions, so the sign on which the CaOx result rests are not specific to one reconstruction.

In the relaxometry-matched formulations the two inversions diverged. MEDI recovered substantially smaller susceptibility magnitudes at the adipose-matched HA locations, −0.128 against −2.683 ppm at 0.70 mm, and detected HA in none of those measurements. The cause is not established. The cause is not established. One possibility is that MEDI's noise weighting suppresses voxels at the particle, which are at 27% of the gel signal. Distinguishing this from other explanations requires further work and is not attempted here.

MEDI required each tube to be inverted separately. Applied to the four tubes together it returned a gel reference standard deviation of 0.219 ppm against 0.007 per tube. Total variation regularised ADMM required no such treatment on the same data.

#### S5. Susceptibility at all particle locations

Streaking is present at the adipose-matched locations, extending several voxels from the particle in a cross pattern, and is absent at the pure alginate and FGT-matched locations reconstructed with the same pipeline. It is not confined to the air-affected locations. The adipose-matched particles are the strongest sources measured (**Table 5**), and streaking from single-orientation dipole inversion scales with source strength.

**Fig. S5.** Susceptibility at all 24 particle locations, at both resolutions, on a common scale. Layout as **Fig. S8**. Negative values are diamagnetic.

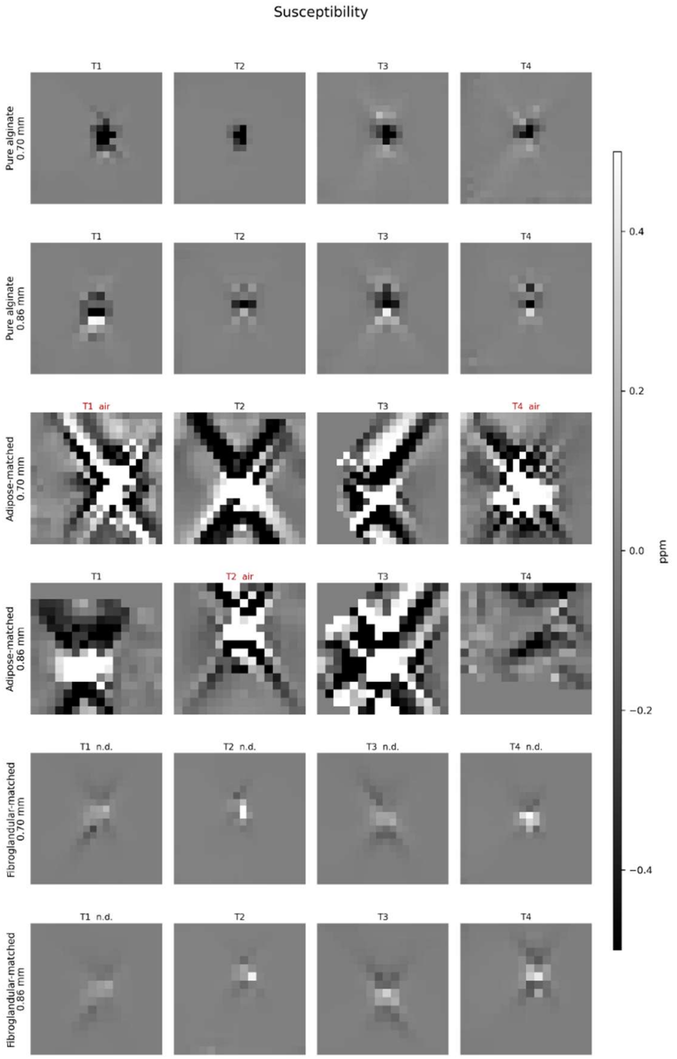

#### S6. All particle measurements

CaOx in pure alginate fails on the focal dip rather than on the threshold. $\Delta\chi_{peak}$ exceeds the threshold in six of eight measurements, but the focal dip is positive in seven of eight: the centre voxel is more paramagnetic than its surroundings, which a diamagnetic source cannot produce.

The threshold criterion alone would therefore report CaOx as detected in six of eight measurements in pure alginate, on susceptibility differences of 0.002 to 0.029 ppm whose centres carry the wrong sign. This is the evidence that the second criterion is necessary rather than redundant.

The same signature appears in FGT HA, where positive focal dips accompany magnitude signal voids. Those locations are independently identified as air-affected by the local field criterion (Section 2.9). The signature is absent from alginate HA, where all eight dips are negative.

**Table S6**. All particle measurements. Δχpeak is the 5th percentile within the two-voxel-radius particle region minus the mean gel reference; the threshold is three times the gel reference standard deviation, measured over a region of the same size. The focal dip is the centre voxel minus the mean of voxels two voxels away; a negative value indicates a localised diamagnetic source. Detection requires both that |Δχpeak| exceeds the threshold and that the focal dip is more negative than one gel standard deviation. The final column names the criterion that failed where detection did not occur.

| Formulation | Res (mm) | Tube | Thresh. (ppm) | HA $\Delta\chi_{peak}$ | HA dip | HA det. | CaOx $\Delta\chi_{peak}$ | CaOx dip | CaOx det. |
|---|---|---|---|---|---|---|---|---|---|
| Alginate | 0.70 | 1 | 0.0060 | -1.4346 | -1.3430 | yes | -0.0167 | +0.0434 | dip |
| Alginate | 0.70 | 2 | 0.0017 | -0.8464 | -1.3639 | yes | -0.0294 | +0.0448 | dip |
| Alginate | 0.70 | 3 | 0.0013 | -0.8559 | -1.5858 | yes | -0.0016 | +0.0828 | dip |
| Alginate | 0.70 | 4 | 0.0044 | -1.2004 | -1.1930 | yes | +0.0066 | +0.0779 | dip |
| Alginate | 0.86 | 1 | 0.0014 | -0.9734 | -0.1207 | yes | -0.0006 | +0.0103 | both |
| Alginate | 0.86 | 2 | 0.0023 | -0.4718 | -0.8603 | yes | +0.0003 | -0.0014 | thresh. |
| Alginate | 0.86 | 3 | 0.0005 | -0.5073 | -1.1147 | yes | -0.0147 | +0.0172 | dip |
| Alginate | 0.86 | 4 | 0.0062 | -0.7430 | -0.5097 | yes | +0.0013 | +0.0045 | both |
| Adipose | 0.70 | 1 | 0.0073 | -2.5029 | -3.0572 | yes | -0.0331 | +0.0423 | dip |
| Adipose | 0.70 | 2 | 0.0287 | -4.8894 | -0.8538 | yes | -0.0866 | +0.0496 | dip |
| Adipose | 0.70 | 3 | 0.0264 | -1.8796 | -0.9810 | yes | -0.0502 | +0.0080 | dip |
| Adipose | 0.70 | 4 | 0.0092 | -1.4607 | -2.9641 | yes | -0.1055 | +0.3059 | dip |
| Adipose | 0.86 | 1 | 0.0080 | -0.7366 | -0.3060 | yes | +0.0104 | +0.0580 | dip |
| Adipose | 0.86 | 2 | 0.0139 | -1.8366 | -2.1246 | yes | +0.0000 | +0.0377 | both |
| Adipose | 0.86 | 3 | 0.0151 | -2.5243 | -1.7062 | yes | -0.4194 | -0.3761 | yes |
| Adipose | 0.86 | 4 | 0.0318 | -0.7579 | -0.6589 | yes | -0.0826 | +0.0345 | dip |
| FGT | 0.70 | 1 | 0.0017 | -0.0360 | +0.1052 | dip | -0.0122 | +0.8021 | dip |

| | | | | | | | | | |
|---|---|---|---|---|---|---|---|---|---|
| FGT | 0.70 | 2 | 0.0158 | -0.0477 | +0.5746 | dip | -0.0601 | -0.0076 | yes |
| FGT | 0.70 | 3 | 0.0062 | -0.0350 | +0.1325 | dip | -1.2757 | -3.0672 | yes |
| FGT | 0.70 | 4 | 0.0040 | -0.0349 | +0.6310 | dip | -0.0049 | -0.0037 | yes |
| FGT | 0.86 | 1 | 0.0063 | -0.0435 | +0.1064 | dip | -0.1099 | +0.9456 | dip |
| FGT | 0.86 | 2 | 0.0036 | -0.0627 | -0.0977 | yes | -0.0281 | +0.1465 | dip |
| FGT | 0.86 | 3 | 0.0003 | -0.1312 | -0.1101 | yes | -0.0008 | +0.0041 | dip |
| FGT | 0.86 | 4 | 0.0052 | -0.1156 | -0.1857 | yes | -0.0882 | +0.8383 | dip |

#### S7. CaOx detection bound

Detection was complete at every value down to −0.10 ppm, partial at −0.05 ppm and absent at −0.02 ppm, placing the limit at approximately 0.05 ppm. HA was detected in all four tubes at every value, confirming that the pipeline behaved identically across the series and that the CaOx outcome reflects the susceptibility supplied rather than a change in the reconstruction.

Recovered values fall consistently below the input, by 30% at −1.82 ppm and 40% at −0.10, which is the dilution described in Section 3.1. The detection limit is therefore a limit on the recovered value rather than on the susceptibility present, and the bound quoted in Section 3.3.3 is expressed on the same basis.

**Table S7**. Detection outcomes in the digital twin across CaOx susceptibility, with HA held at −2.0 ppm as an internal control. [n = 4 tubes per condition.]

| $\chi_{CaOx}$ (ppm) | recovered peak (ppm) | CaOx detected | HA detected |
|---|---|---|---|
| −1.82 | −1.274 | 4/4 | 4/4 |
| −1.41 | −1.004 | 4/4 | 4/4 |
| −1.00 | −0.807 | 4/4 | 4/4 |
| −0.50 | −0.395 | 4/4 | 4/4 |
| −0.25 | −0.185 | 4/4 | 4/4 |
| −0.10 | −0.065 | 4/4 | 4/4 |
| −0.05 | −0.030 | 2/4 | 4/4 |
| −0.02 | −0.012 | 0/4 | 4/4 |

#### S8. R2* maps at all particle locations

Both mineral types elevate R2* relative to the surrounding gel, at every location and both resolutions, in contrast to the susceptibility maps of Fig. S8 in which only HA produces a focal source. The difference in baseline between formulations is apparent: the adipose-matched gel is an order of magnitude higher than pure alginate, so the particle contribution sits on a larger and more variable background.

**Fig. S8**. R2* at all 24 particle locations, at both resolutions. Rows are formulation and resolution; columns are the four tubes, HA first and then CaOx. All panels share one scale. Locations identified as air-affected (**Section 3.5.1**) and those not meeting the detection criterion are labelled. Grey indicates voxels outside the mask. Panels differ in physical extent between the two resolutions.

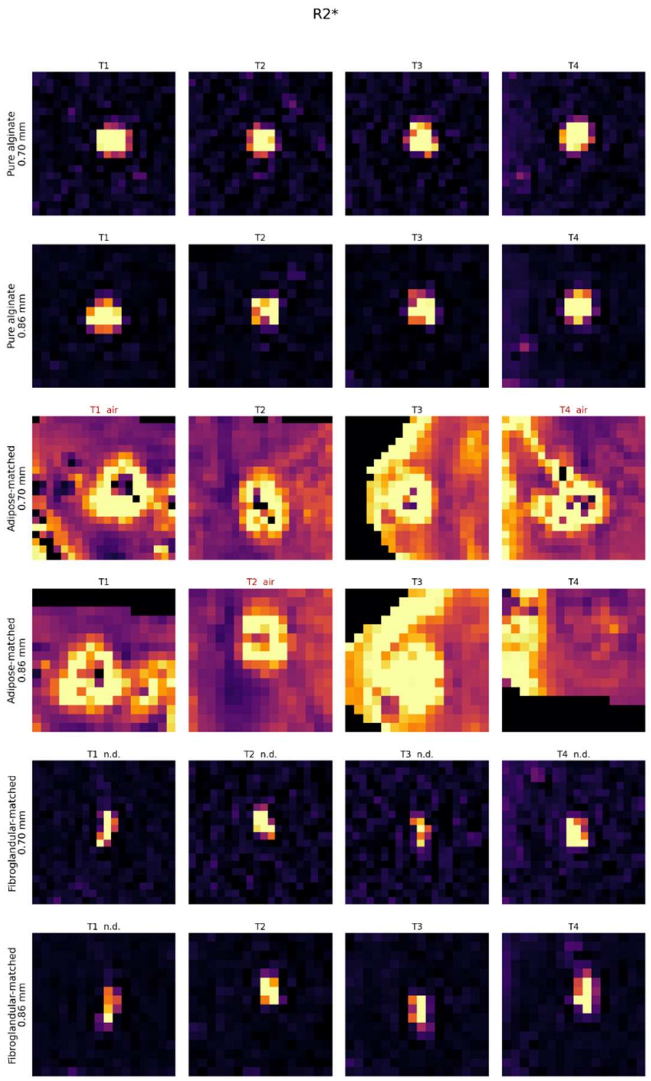
